# Key link between iron and the size structure of three main mesoplanktonic groups (Crustaceans, Rhizarians, and colonial N_2_-fixers) in the Upper Ocean

**DOI:** 10.1101/2024.03.08.584097

**Authors:** Mathilde Dugenne, Marco Corrales-Ugalde, Jessica Y. Luo, Lars Stemmann, Jean-Olivier Irisson, Fabien Lombard, Todd O’Brien, Charles Stock, PSSdb data contributors, Rainer Kiko

## Abstract

In marine ecosystems, critical services like fish production, carbon export, or the delivery of nutrients through N_2_-fixation rely heavily on the size spectrum of pelagic organisms, particularly mesoplankton (200-20,000 µm). However, how environmental factors shape mesoplankton spectral biogeography remains largely unresolved, as so far only limited datasets exist to understand the large-scale shifts in mesoplankton size. Using global compilations of Rhizarian, colonial N_2_-fixer, and Crustacean images, we reveal the paramount role of iron in shaping the size structure and related biogeography of these groups. Our findings underscore the importance of atmospheric sources of iron for N_2_-fixers and Rhizarians while total iron, accounting for organic and inorganic compounds, appeared to explain most of the variance in Crustacean size structure via apparent recycling. With a comprehensive set of explanatory variables, our models reached high R^2^ (0.93, 0.61, and 0.69 respectively), providing robust predictions of mesoplankton size structure related to elemental cycling and ecosystem services. Our results suggest that future increases in global temperatures will have negative effects on mesoplankton size, possibly limiting carbon export from the productive layers to sequestration depth, that can be offset by expected increases in iron inputs that benefit N_2_-fixers, Rhizarians, and eventually Crustaceans.

## Introduction

The decrease of organism abundances in increasing size classes is a universal feature of both terrestrial and aquatic ecosystems (^1–4^) that relates to ecosystem functioning and services like carbon export and fish biomass production for the latter (^5-8)^. Both services are affected by size-dependent processes, such as CO_2_ uptake (^9^) and respiration (^10^), predation (^11^), organic matter packaging (^12,13^) and passive sinking (^14^), that underpin all links of marine food webs, from bacteria to whales (^15-17)^. As a result, changes affecting plankton communities are currently observed through the lens of the size spectrum framework via three parameters (^18^): the spectral intercept (which will increase or decrease if all plankton groups are equally affected), the slope (which will vary with differences in transfer efficiency), and the linearity (which may decrease under perturbations that alter only certain taxa). Using this framework, ecologists have shown that most plankton communities varied in size with latitudinal gradients (^19,20^), following the set of empirical rules and theoretical principles linking environmental gradients to plankton size structure reviewed by Barton et al. (^21^). In their review, the authors identified seasonal and latitudinal variations of temperature and oxygen levels, resource availability and prey size structure, as well as ocean circulation as major structuring elements of the pelagic realm. Yet in practice, their effects are often hard to disentangle as most factors co-vary in marine ecosystems and some relationships appear contradicting to explain the spectral biogeography of the three groups that dominate mesoplankton communities in the upper ocean (^22^): heterotrophic Crustaceans, mixotrophic Rhizarians, and autotrophic N_2_-fixers. For example, Crustacean body size decreases under high temperature and low oxygen levels, a relationship known as the Temperature-Size Rule (^23^), but also surprisingly with phytoplankton productivity and cell size (^24^). Opportunist mixotrophs are equally unpredictable, as they should be favored in oligotrophic ecosystems, where sources of inorganic and organic nutrients are scarce, and paradoxically in productive systems that undergo large seasonal fluctuations in resource availability (^21^). Lastly, future projections of marine N_2_-fixation are still difficult despite its importance in delivering new nutrients to oligotrophic ecosystems (^25^), as greater temperatures should increase the growth of large N_2_-fixers like *Trichodesmium*, but also limit the supply of their essential nutrients that would require lower surface:volume ratio. Overall, previous meta-analysis presenting the biogeography of crustacean copepod (^24^), mixotrophic (^26^) and heterotrophic Rhizarians (^27^), and colonial N_2_-fixers (^19^) only explained up to 50% of the variance in mesoplankton occurrences or sizes, leaving important sources of variability unexplained that the authors often linked to missing explanatory variables or to further clustering within broad groups. Eventually, all relationships linking environmental factors to organism size spectra remain largely enigmatic, in part due to the paucity of direct size structure compilations and related environmental gradients, and to additional traits (diet, morphotypes) underpinning specific adaptations within broader phyla.

In recent years, the size-structure framework has been used in conjunction with other traits (e.g. feeding strategy, migration, buoyancy) to represent plankton communities along a few functional axes, instead of numerous taxa that cannot all be monitored or cultivated (^28-35^). To do so, ecologists have been using new imaging devices that provide high-throughput proxies of functional complexity across the five orders of magnitude in plankton sizes (^36,37^). Here, we present a compilation of keystone mesoplankton groups size spectra measured from imaging devices deployed globally between 0-200m. We depict the spectral biogeography of Crustaceans based on net collected samples imaged with benchtop scanners, as well as Colonial N_2_-fixers and Rhizarians using Underwater Vision Profilers (UVP), since these organisms are severely damaged by nets. We first explore the relationships between their size distributions and environmental descriptors (e.g. temperature, resource availability, or mesoscale circulation), including diverse iron supply proxies whose importance in explaining mesoplankton biogeography will be tested for the first time, before searching for specific adaptations within these broad groups. Ultimately, we aim at improving our mechanistic understanding of marine ecosystem functioning under diverse stressors, as we used the linkages between size structure and environmental gradients in conjunction with machine learning models (i.e. boosted regression trees) to generate global predictions of group-specific size structure. These will help refine or validate plankton models, with the goal to constrain the future projections of ecosystem processes under climate change.

## Results

### A spectral analysis of three main mesoplankton biogeography in the Upper Ocean

In this study, we compiled images from benchtop scanners and UVP that have been regularly deployed in all major ocean basins since the mid 2000’s, with the highest concentration of samples recorded within the epipelagic layer of the subtropics (Figure 1a-b). Despite their relative spatial proximity, regions with the strongest sampling bias were typically sampled months apart (Figure 1c-f), providing discontinuous records of mesoplankton size spectrum in space and time that helped us disentangle the effect of seasonal or decadal trends from broad spatial gradients on their spectral biogeography. After taxonomic standardization, all image annotations pointed to the global importance of colonial N_2_-fixers, Rhizarians and Crustaceans within mesoplankton communities, as the first two groups represented 45% and 15% of the total images compiled from UVP datasets, while the latter corresponded to 81% of the overall scanner images aligning with the known bias for non-gelatinous taxa collected with traditional nets (Figure 1g-h). We combined UVP profiles and scanned nets to compute the epipelagic size spectra of colonial N_2_-fixers (exclusively represented by two morphotypes of the colonial genus *Trichodesmium*), Rhizarians (regrouping the 4 main classes of Acantharia, Collodaria, Foraminifera, and Phaeodaria), and Crustaceans (including three classes of Copepoda, Malacostraca, and Ostracoda), averaged by 1° latitude longitude and by year month bins to limit statistical bias arising from auto-correlated measurements.

**Figure 1.**
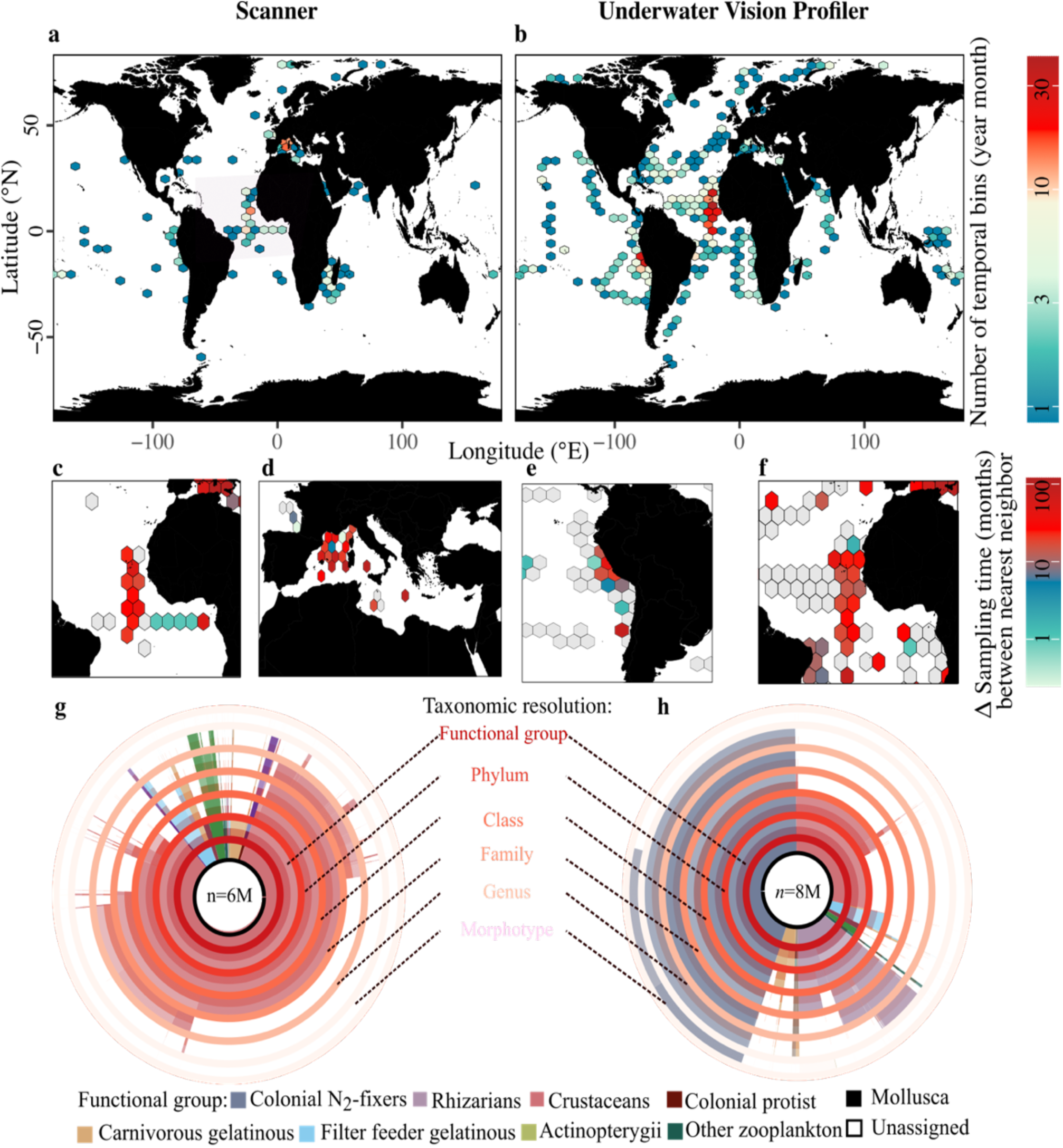
Resolution and diversity in taxonomic annotations of the Scanner (left) and UVP (right) imaging datasets. Spatial coverage of the scanner (**a)** and UVP (**b)** datasets produced in the Pelagic Size Structure database. The colors represent the number of temporal (year month) bins in these datasets. Each spatial bin has been sampled irregularly over time, often months apart in the most biased regions of the tropical Atlantic (**c & f**), Mediterranean Sea (**d**), or eastern Equatorial Pacific (**e**). Hence our datasets represent a unique opportunity to study how mesoplankton size structure fluctuate in space and time because of environmental variables. Diversity of taxonomic annotations (*n* represent the total number of images in each dataset) separated by taxonomic ranks (red to pink rings) and colored by plankton functional groups in the scanner (**g**) and UVP (**h**) datasets. Blank portions correspond to organisms unassigned at the considered ranks.

Important similarities across the three main groups’ size spectra can be seen in the global compilation (Figure 2a), as they all show a log-linear decline in abundance with increasing size classes following slopes close to the canonical value of -1 (Supplementary Figure 1). Yet, their total abundances, obtained by integrating individual size spectra across all observed sizes, span four orders of magnitude, ranging between ∼0.1 and ∼1000 particles m^-3^. This represents a significant spatio-temporal variability across the PSSdb datasets, particularly for colonial N_2_-fixers which present the highest percentage of empty records, with 64% of the spatio-temporal bins without occurrences (Figure 2b), compared to Rhizarians (36%) and Crustaceans (12%).

**Figure 2.**
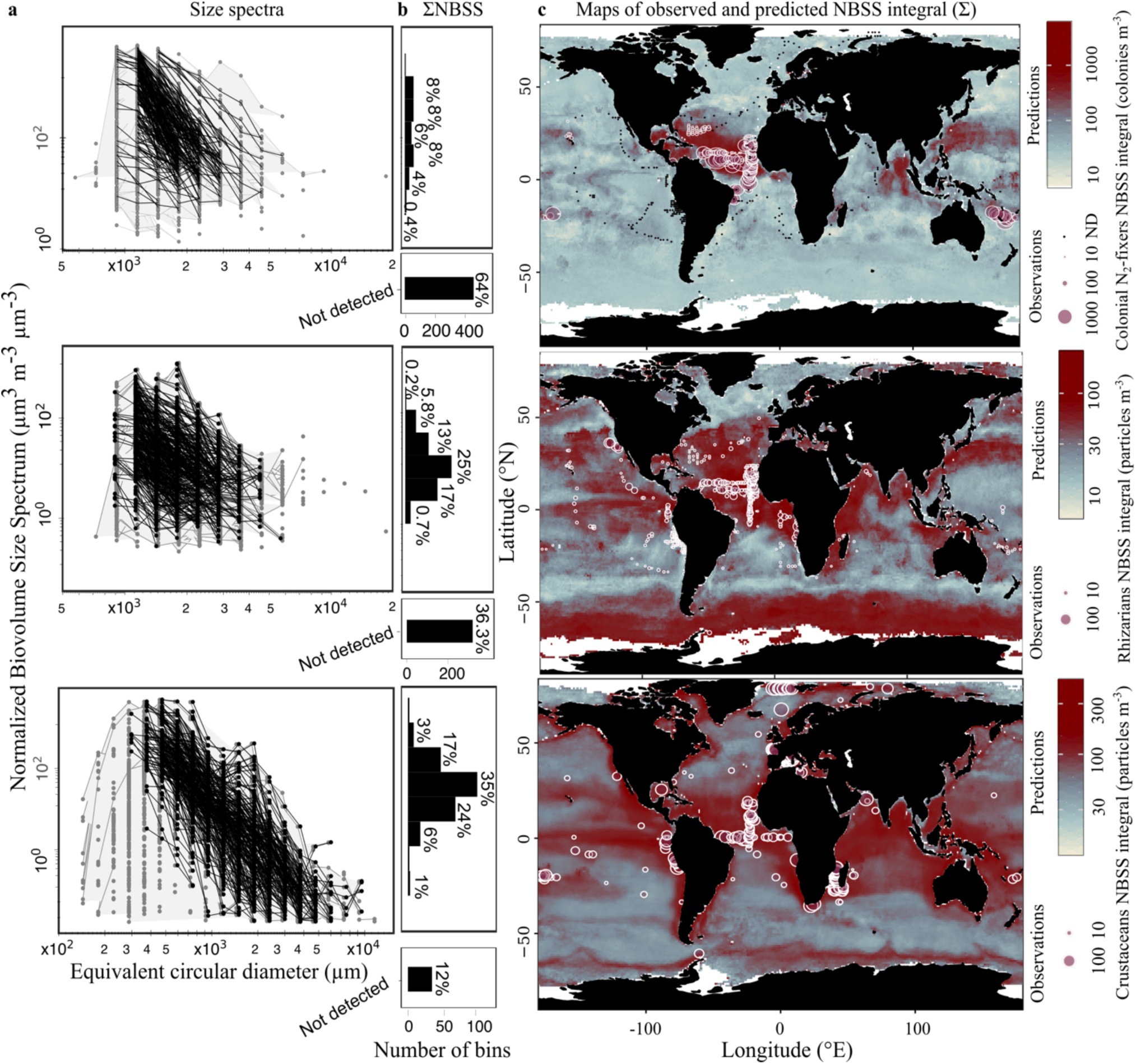
Observed size spectra (left) and maps of observations and predictions (right) of colonial N_2_-fixers (top panels), Rhizarians (middle panels), and Crustaceans (bottom panels) total abundance. **a** Group-specific size spectra derived from UVP (for colonial N_2_ fixers on top and Rhizarians in the middle) and scanner (for Crustaceans at the bottom) images. Gray dots mark size classes that were filtered out due to methodological bias (mesh of the nets, particles too rare to be counted accurately) and black dots indicate all final data points used to infer spectral parameters and predict NBSS. Note that x-axis limits differ depending on the instrument resolution and detection limits. **b** Total abundance of colonial N_2_ fixers (top), Rhizarians (middle), and Crustaceans (bottom) in individual grid cells (1°x1° longitude/latitude and year month) distributed globally. Note that the y-axis is shared for panels **a** & **b**. Number and percentage of empty records (not detected) are also indicated in separate sub-panels beneath each row. **c** Maps of observed (open circles) and predicted (colored tiles) average abundance, obtained by integrating NBSS estimates using predicted parameters (slope, intercept, size range). ND: Not detected.

We found that colonial N_2_-fixers were largely restricted to the subtropical latitudes, between -20 and 20 °N, similarly to Rhizarians though they were also relatively abundant at higher latitudes and in the Californian upwelling system (Figure 2c).

In comparison, Crustaceans were generally ubiquitous, with the highest abundances observed in productive ecosystems (e.g. polar, upwelling and coastal regions). Caution should be taken when using or interpreting the few observations located in the Southern Ocean, but our results shed lights on previously undersampled regions that could be potential hotspots for marine mesoplankton, as reflected in other parts of the globe where environmental conditions are highly similar (Arctic Ocean, upwellings, etc.).

### Iron, a key descriptor of mesoplankton spectral biogeography in the Upper Ocean

Boosted regression tree models fitted to PSSdb records (response variable) and global datasets of environmental descriptors (explanatory variables) yielded global predictions that match the magnitude and variability of total abundances derived from spectral parameters (Figure 2c) with relatively high R^2^ (0.93, 0.61 and 0.69 for N_2_-fixers, Rhizarians and Crustaceans respectively, Table 1). The main bottom-up factors explaining the spectral biogeography of Crustaceans, Rhizarians and colonial N_2_-fixers, indicated by their respective ranks in terms of their gradual importance in the model variance explained, typically included temperature as well as diverse iron proxies (Table 1).

**Table 1.** Goodness of fit of mesoplankton size spectra predicted from a boosted regression tree model tested with the full set of environmental factors or with a reduced model excluding all iron proxies. Ranking of environmental variables in the full model expressing their importance in variance explained (in parenthesis).

| Model | Colonial $N_2$-fixers | | Rhizarians | | Crustaceans | |
| --- | --- | --- | --- | --- | --- | --- |
|  | Full | Reduced | Full | Reduced | Full | Reduced |
| $R^2$ | 0.93 | 0.61 | 0.61 | 0.38 | 0.69 | 0.35 |
| Rank in importance: |  |  | Variables (feature importance metric): |  |  |  |
| 1 | Temperature (0.39) |  | Temperature (0.53) |  | Total iron (0.15) |  |
| 2 | Phosphates (0.18) |  | Wet dust deposition (0.08) |  | Photosynthetic Available Radiance (0.11) |  |
| 3 | Aerosol (0.13) |  | Apparent Oxygen Utilization (0.06) |  | Sea Level Anomaly (0.11) |  |
| 4 | Total iron (0.11) |  | Photosynthetic Available Radiance (0.06) |  | Temperature (0.08) |  |
| 5 | Wet dust deposition (0.09) |  | Total iron (0.06) |  | Mixed Layer Depth (0.08) |  |
| 6 | Photosynthetic Available Radiance (0.04) |  | Finite Size Lyapunov Exponent (0.06) |  | Particle Size Distribution slope (0.07) |  |
| 7 | Mixed Layer Depth (0.04) |  | Mixed Layer Depth (0.05) |  | Chlorophyll concentration (0.05) |  |
| 8 | Apparent Oxygen Utilization (0.04) |  | Sea Level Anomaly (0.05) |  | Apparent Oxygen Utilization (0.05) |  |
| 9 | Silicates (0.04) |  | Phosphates (0.05) |  | Aerosol (0.04) |  |
| 10 | Particle Size Distribution slope (0.04) |  | Particle Size Distribution slope (0.05) |  | Phosphates (0.04) |  |
| 11 | Nitrates (0.04) |  | Nitrates (0.05) |  | Finite Size Lyapunov Exponent (0.04) |  |
| 12 | Fraction of picophytoplankton (0.02) |  | Chlorophyll concentration (0.04) |  | Wet dust deposition (0.03) |  |
| 13 | Fraction of Microphytoplankton (0.02) |  | Silicates (0.03) |  | Nitrates (0.03) |  |
| 14 | Sea Level Anomaly (0.02) |  | Fraction of picophytoplankton (0.03) |  | Fraction of picophytoplankton (0.03) |  |
| 15 | Chlorophyll concentration (0.02) |  | Fraction of nanophytoplankton (0.03) |  | Nanophytoplankton (0.03) |  |
| 16 | Finite Size Lyapunov Exponent (0.02) |  | Aerosol (0.03) |  | Silicates (0.02) |  |
| 17 | Fraction of nanophytoplankton (0.02) |  | Fraction of microphytoplankton (0.03) |  | Fraction of microphytoplankton (0.02) |  |

Our full model included several proxies linked to different, but not mutually exclusive, pools of iron to reflect the diversity in natural iron compounds found in marine ecosystems (^38^). Labile compounds were mostly represented by aerosols, either from desert sources (best approximated by wet dust deposition satellite products) or from either volcanic and wildfire ashes or anthropogenic activities (all affecting the total aerosol optical thickness satellite product). A third proxy, representing the total iron concentration accounting for both mineral and organically-bound Fe ligands, from the biogeochemical model PISCES, was also tested given the similarity found between modeled concentrations and monthly average observations of *in situ* dissolved iron concentration measured as part of the GEOTRACES program (Supplementary Figure 2). When iron proxies were excluded in our model construction, the total variance explained was reduced by 40-50% in the case of Rhizarians and Crustaceans, for which dust deposition and total iron ranked second and first respectively, and by a third for colonial N_2_-fixers, whose biogeography appeared more linked to aerosol emissions alongside temperature and phosphate concentration (Table 1).

The temporal correlation between predicted abundances and the different iron proxies in individual grid cells (Supplementary Figure 3) highlights the importance of mineral and other organically-bound iron proxies in mesoplankton model predictions (Figure 3). The seasonal trends of NBSS predictions shows important similarities with dust and aerosol proxies, ranked 5^th^ and 3^rd^ in model’s feature importance, suggesting a key role in shaping the size distribution of N_2_-fixers in the Pacific and Atlantic subtropical gyres respectively (Figure 3a-b, top panels). In the former ecosystem, a main peak in abundance is observed in winter (Oct-Dec in the Northern hemisphere and Jun-Aug in the Southern hemisphere), consistent with a negative correlation between N_2_-fixer abundances and light availability (Supplementary Table 1) and relatively high aerosol emissions (Figure 3b). Correlations with aerosols were highest in the South Pacific Subtropical Gyre (SPSG), where anthropogenic dust is supplemented by ashes produced by volcanic activities or wildfires. In the Atlantic, where the main iron pool is provided by Saharan dust deposition, seasonal trends of N_2_-fixer NBSS strongly followed the climatology of the dust product, marked by a peak of iron supply early in the year that progressively decreases to a minimum observed in June during the dry season. Similarly, N_2_-fixer abundances were low until June and increased later in the year.

**Figure 3.**
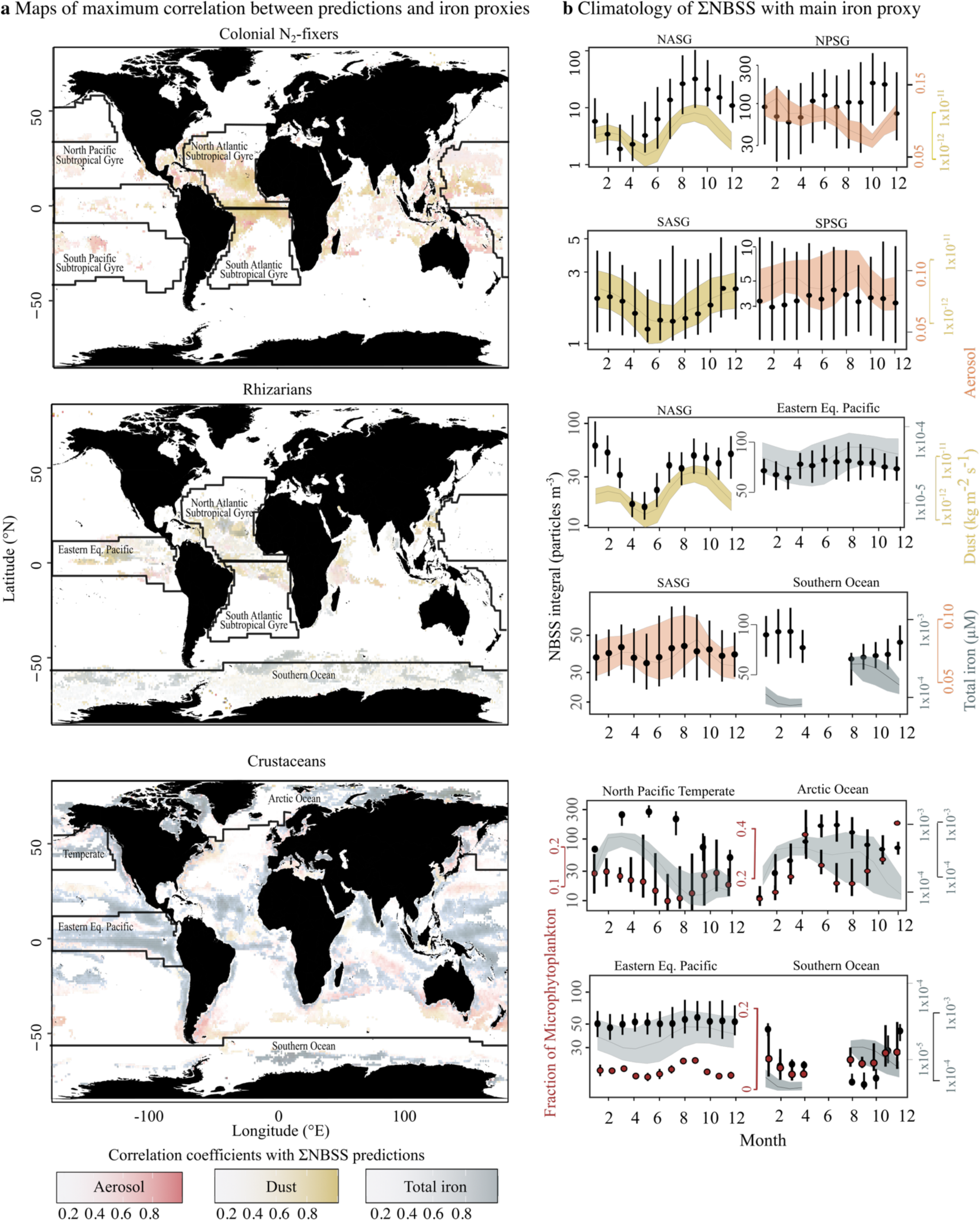
Main iron proxies linked to the spectral biogeography of colonial N_2_-fixers (top panels), Rhizarians (middle panels), and Crustaceans (bottom panels). **a** Maps of maximum correlation coefficients between predicted NBSS integral and multiple iron proxies used to determine the main proxy in specific ecosystems. The main iron proxy was attributed according to the variable counting the most grid cells in specific ecosystems and ocean regions. Ecosystems whose climatology is presented in panel c are outlined in black. **b** Climatology of mesoplankton groups (black dots) and the main iron proxy (ribbons) in ecosystems with significant correlations. Seasonal trends of microphytoplankton (red dots) are superimposed on the climatology of Crustaceans NBSS integral to link their dynamics to that of phytoplankton prey.

Likewise, both aerosols and wet dust deposition seemed to be driving the relatively high abundance of Rhizarians in most of the subtropical gyres (Figure 3a-b, middle panels). The importance of aerosols was marked in several basins near the Equator, including in the Indian, Pacific and Atlantic oceans. Observed correlation with specific classes of Rhizarians indicated that while aerosols were positively related to all classes, only Collodarians seemed to be favored by dust deposition (r=0.28, Supplementary Table 1) with a peak in abundances marked after the input of Saharan dust during the wet season late in the year. Interestingly, the peak of modeled abundance of Rhizarians (Figure 2c) in the Southern Ocean and upwelling ecosystems was likely linked to increased concentrations of total iron (Figure 3a-b), as monthly average observations of *in situ* dissolved iron concentration measured as part of the GEOTRACES program also matched that of Rhizarian groups (i.e Phaeodaria and Foraminifera), with very high correlation coefficients (r>0.77, Supplementary Table 1). This gives us further confidence in the linkages found between the total iron model product and mesoplankton spectral biogeography.

Compared to N_2_-fixers and Rhizarians, Crustaceans were almost exclusively linked to total iron concentration in all ecosystems (Figure 3a-b, bottom panels). This result was also evident from the correlation with *in situ* observations from the GEOTRACES program, especially for copepods with respective coefficients of 0.44 with PISCES iron and 0.5 with GEOTRACES iron (p-value<0.001, Supplementary Table 1). As a result, predictions of Crustacean total abundance closely followed the seasonal trends in total iron concentrations, especially in productive regions like temperate, polar, coastal, or upwelling ecosystems (Figure 2c, 3c). In the North Pacific Temperate latitudes, Crustacean abundances increased in the spring, following the peak in microphytoplankton size fraction observed by satellite. Likewise, the pool of total iron presented a peak of concentration between Feb-Apr, following the increased Ekman pumping and deep mixing events occurring in the wintertime. In comparison, the wind-driven Ekman pumping located in the Eastern Equatorial Pacific is more sustained throughout the year (^39^), although Trade winds generally intensify between June-August, resulting in a small increase in total iron and overall weaker seasonal trends caused by upwelling (Figure 3c). Yet, both the microphytoplankton fraction observed by satellite and our predictions of Crustacean abundances seemed to be linked to the overall availability of total iron in this ecosystem, with a concentration peak recorded in September. In polar ecosystems, our knowledge on seasonal trends is limited by the sea ice coverage, the lack of sunlight and higher cloud coverage in the wintertime, as highlighted in important temporal and spatial gaps in both satellite and model products (Supplementary Figure 2), resulting in only partial predictions of Crustaceans size distribution at least in the Southern Ocean (Figure 2 & 3). Despite such gaps, it appeared that total iron concentration and Crustacean abundances were higher during the austral summer than during the winter (Figure 3c), which is unexpected given that phytoplankton blooms initiated in the summer are thought to considerably drawdown surface iron concentrations (^40^). This phenomenon was even more apparent in the Arctic Ocean where the demise of the spring bloom does not appear to cause a decline in both iron and Crustacean concentrations, which both remain relatively high through the end of year.

### Adaptations and partitioning of spectral biogeography within broad mesoplankton groups in the Upper Ocean

Grouping image annotations by finer taxonomic resolution (classes in the case of Rhizarians and Crustaceans) or by morphotypes (in the case of colonial N_2_-fixers) resulted in overall improved model performances compared to the full model presented above (Table 2), except for rare groups of Phaeodaria (R^2^=0.26) and Foraminifera (R^2^=0.25). The decrease in model information criteria (BIC and AIC estimates), indicative of improved fit, were more marginal for the colonial N_2_-fixers due to the dominance of a specific class (i.e. Oscillatoriales) and morphotype (i.e. Tuff) in the UVP images (data not shown). In comparison, both Rhizarians and Crustaceans presented larger differences in model performance metrics when images were grouped in higher taxonomic ranks (classes or families) despite lower available observations, supporting strong evidence for their niche partitioning in the upper ocean. Using the same modeling approach applied to finer taxonomic groups, including copepods specific families that often correlate with a certain diet (e.g. omnivores or carnivores) as compiled in Benedetti et al. (^41^), we highlight the potential adaptations or plasticity, in the case of *Trichodesmium* morphotypes since multiple species can form either morphotype, of these subgroups to certain environmental conditions.

**Table 2.** Statistical metrics to evaluate the goodness of fit of mesoplankton size spectra predictions in increasing taxonomic ranks. Optimal models are indicated by low AIC/BIC and high R^2^

| Taxa | Colonial *N*<sub>2</sub>-fixers |  |  | Rhizarians |  | Crustaceans |  |  |
| --- | --- | --- | --- | --- | --- | --- | --- | --- |
|  | Phylum | Family | FG | Phylum | Class | Phylum | Class | FG |
|  | Cyanobacteria | Oscillatoriales | Puff<br>Tuff | Radiozoa | Acantharia<br>Collodaria<br>Phaeodaria<br>Foraminifera | Crustacea | Copepoda<br>Malacostraca<br>Ostracoda | Carnivore<br>Omnivore |
| BIC | 145.92 | 145.4 | 143.2 | 150.5 | 132.7 | 145.6 | 135.9 | 134.3 |
| AIC | 144.1 | 144.1 | 142.1 | 149.1 | 133.4 | 152.2 | 134.7 | 135.1 |
| R<sup>2</sup> | 0.93 | 0.93 | 0.93<br>0.95 | 0.61 | 0.54<br>0.66<br>0.26<br>0.25 | 0.69 | 0.84<br>0.62<br>0.43 | 0.68<br>0.62 |

Functional groups (FG): Morphotypes for colonial *N*<sub>2</sub>-fixers or trophic group for copepods according to the compilation of Benedetti et al. <sup>(41)</sup>

To identify the environmental factors involved in subgroup adaptations and assess the degree of partitioning within subgroups represented in Figures 4-6a, we used the partial dependency responses of both spectral intercept and slope to all environmental factors tested. Partitioning was then calculated as the portion of the area under the curve that was not shared amongst all subgroups, which can be compared across environmental descriptors, and also across broad mesoplankton phyla. Many iron proxies were involved in niche partitioning, indicating that the link between iron proxies and mesoplankton biogeography discussed in the section above was often caused by one, often dominant, taxon. For example, aerosol thickness seemed to promote both the intercept and slope of Puff (r=0.14 and 0.22 respectively, Supplementary Table 1 and Figure 4b) rather than Tuff colonies, which better responded to phosphate concentration (r=-0.28 and 0.22 respectively, Supplementary Table 1 and Figure 4b). This distinction caused the former to be more abundant in regions where aerosols are released from hydrothermal vents near the Tonga islands and anthropogenic activities located in Asia in the Pacific Subtropical Gyres (Figure 4c). Similarly, we found that both aerosol and dust emissions must contribute significantly to the hotspots of two mixotrophic Rhizarian classes, Collodarians and Acantharians, in the Atlantic and Pacific Gyres (Figure 5c). Inversely, heterotrophic members like Phaeodaria and Foraminifera seemed more adapted to total iron concentrations, promoting larger colonies (r=0.24 and 0.18 respectively, Supplementary Table 1 and Figure 5b) that also benefited from higher chlorophyll-a concentration (r=0.26 respectively, Supplementary Table 1 and Figure 5b). As a result, these classes were dominant in productive systems located in polar regions, near the coasts, and in the upwelling system off the Californian coast (Figure 5c). Total iron concentrations were also important in driving Crustaceans adaptations, as mostly Copepoda appeared correlated to increased trace metal availability, underlying the excessive dominance of Copepoda, especially omnivorous families, in polar and other productive ecosystems. Conversely, Ostracoda seemed more adapted to stratified ecosystems (r(Sea Surface Temperature, NBSS intercept)=0.41 and r(Mixed Layer Depth, NBSS intercept)=-0.07), Supplementary Table 1) located in the subtropics, while Malacostraca appeared to strive under higher nitrate concentrations (r=0.1) (Figure 6).

**Figure 4.**
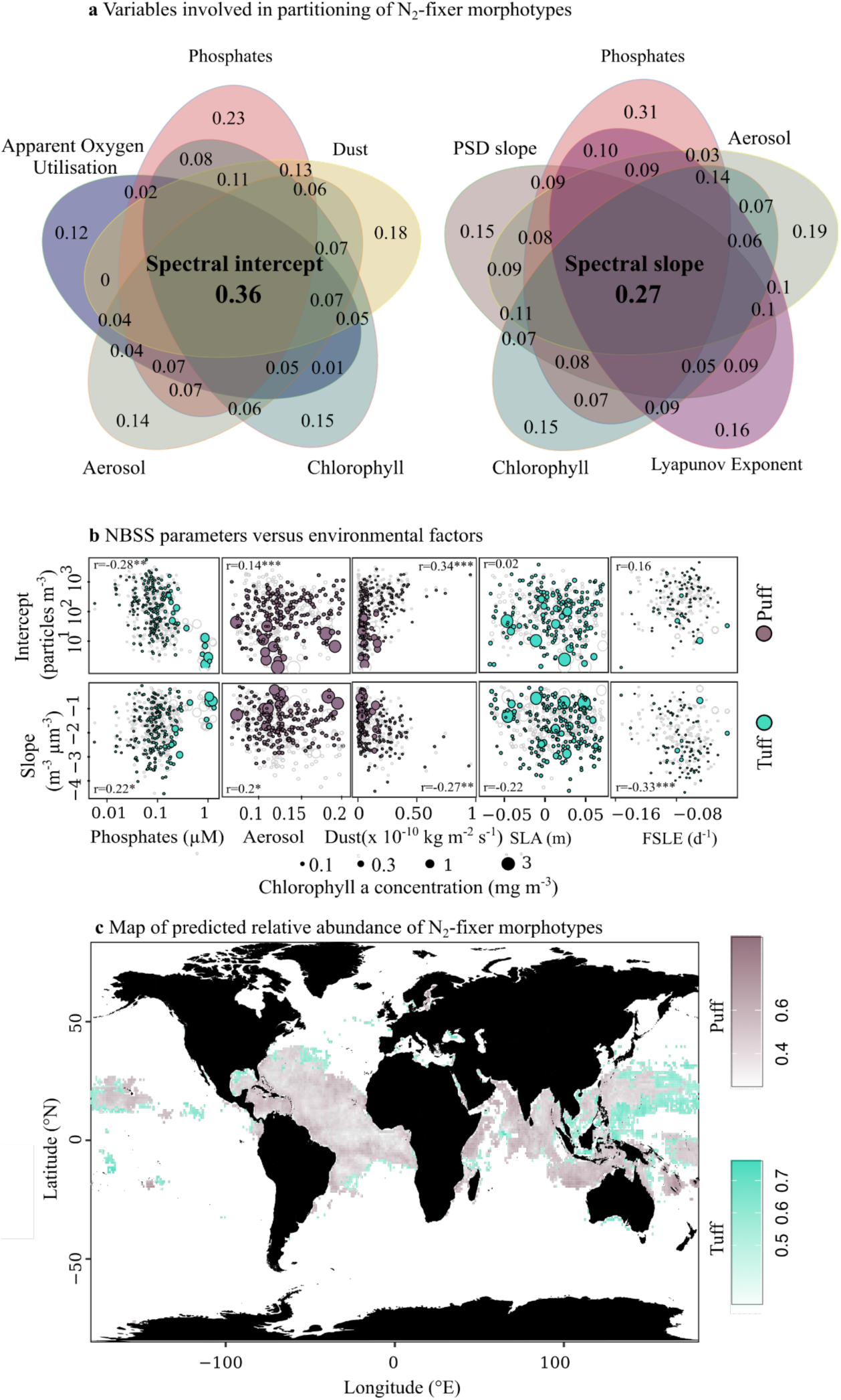
Environmental plasticity of colonial N_2_-fixers morphotypes. **a** Venn diagram representing the top-5 stressors involved in morphotype partitioning (numbers represent the degree of partitioning) based on partial dependence plots for spectral intercept (left diagram) and slope (right diagram) **b** Correlations between environmental factors and Tuff (green dots) or Puff (brown dots) NBSS parameters (slope and intercept). Datapoints vary in size according to the mean chlorophyll-a concentration in that same bin. **c** Predicted maps of colonial N_2_-fixer morphotypes relative abundance (same colorcode). SLA: sea level anomaly, FSLE: finite size Lyapunov exponent

**Figure 5.**
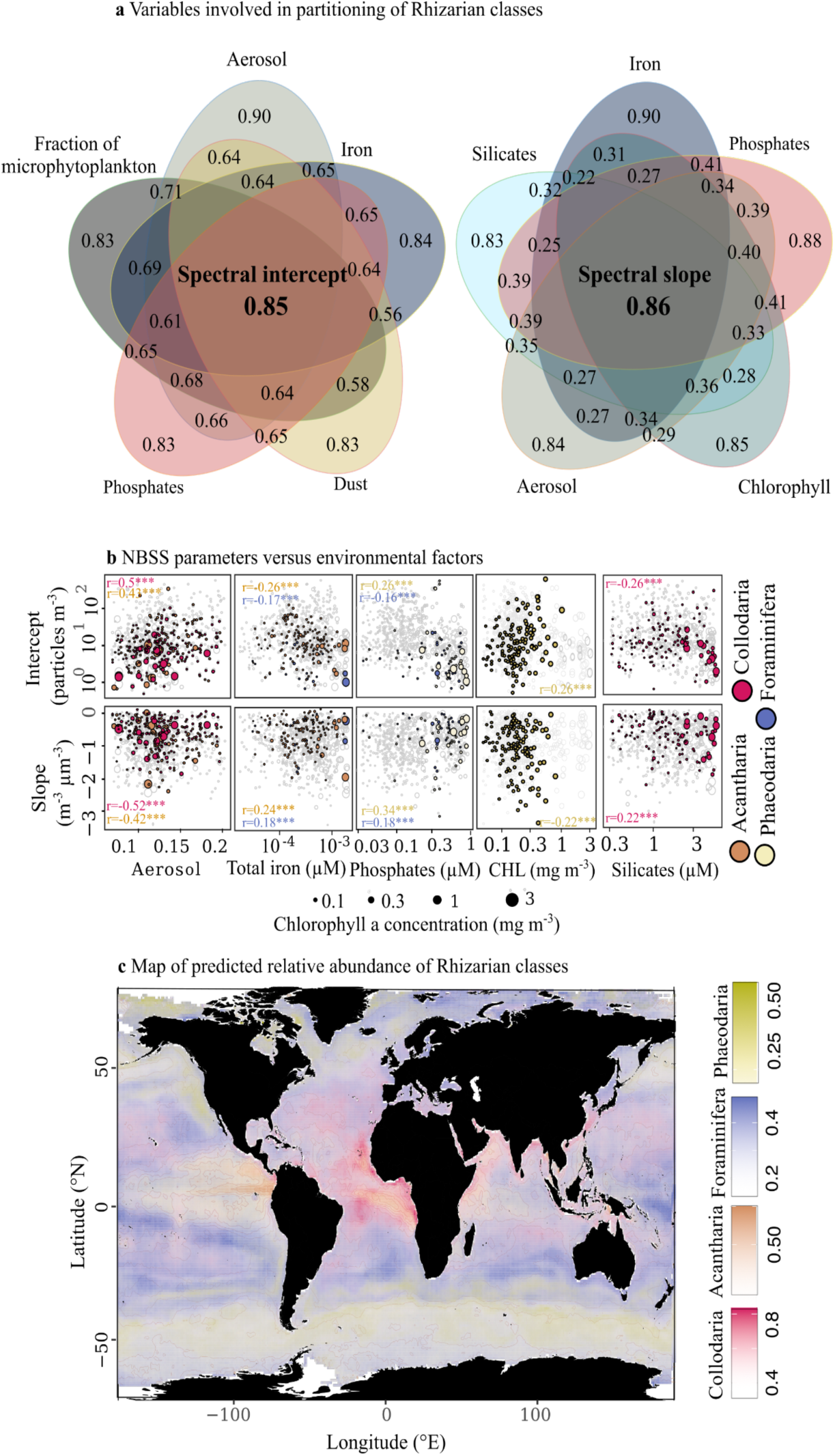
Environmental adaptations within Rhizarian classes. **a** Venn diagram representing the top-5 stressors involved in classes partitioning (numbers represent the degree of partitioning) based on partial dependence plots for spectral intercept (left diagram) and slope (right diagram) **b** Correlations between environmental factors and Acantharia (orange), Collodaria (pink), Foraminifera (blue), or Phaeodaria(yellow) NBSS parameters (slope and intercept). Datapoints vary in size according to the mean chlorophyll-a concentration in that same bin. **c** Predicted maps of Rhizarian classes relative abundance (same colorcode). CHL: surface chlorophyll-a concentration

**Figure 6.**
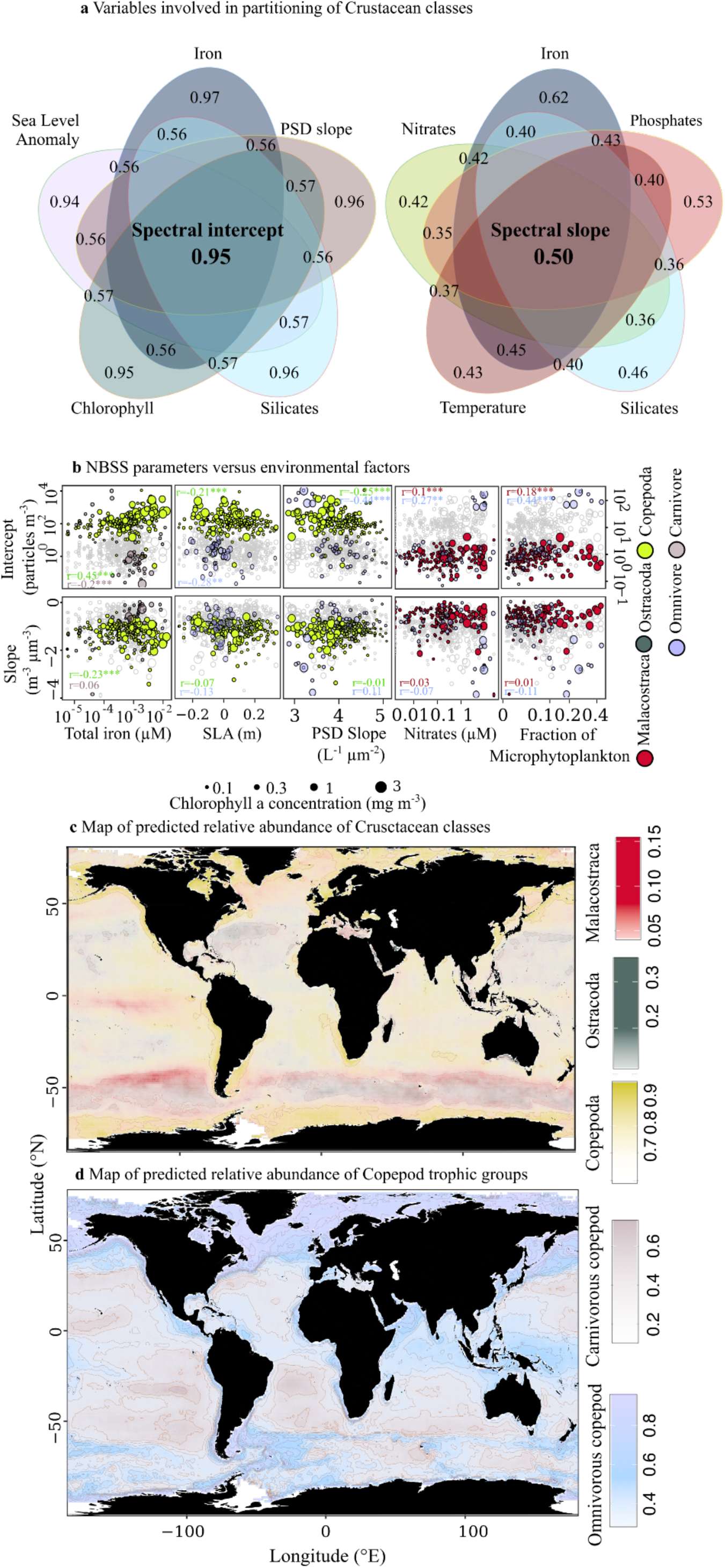
Environmental adaptations within Crustacean classes and trophic groups. **a** Venn diagram representing the top-5 stressors involved in classes partitioning (numbers represent the degree of partitioning) based on partial dependence plots for spectral intercept (left diagram) and slope (right diagram) **b** Correlations between environmental factors and Copepoda (yellow), Ostracoda (green), Malacostraca (red), and copepod functional groups (omnivore in blue carnivore in grey) NBSS parameters (slope and intercept). Datapoints vary in size according to the mean chlorophyll-a concentration in that same bin. **c** Predicted maps of Crustacean classes relative abundance (same colorcode). SLA: sea level anomaly, PSD slope: Slope of the Particle Size Distribution

Examples of second order proxies involved in niche partitioning also included indicators of mesoscale circulation and availability of other nutrients, namely phosphates for colonial N_2_-fixers and Foraminifera, silica for siliceous mixotrophic Rhizarians (e.g. Collodaria) and omnivorous copepods, or large prey (indicated by chlorophyll-a concentration, higher fraction of microphytoplankton, and lower absolute slope of the particle size distribution) for flux-feeding Phaeodaria, omnivorous copepods, and Malacostraca (Supplementary Table 1 and Figures 4-6). By pairing class- or morphotype-specific spectral biogeography to mesoscale circulation proxies, we found that elevated concentrations of both Copepoda and Ostracoda were linked to cyclonic eddies (Figure 6b), indicated by negative sea level anomalies (r=-0.2 and -0.16 respectively, Supplementary Table 1), as opposed to the class Malacostraca. While SLA did not impact the intercept of colonial N_2_-fixers NBSS significantly, their slopes appeared relatively flatter in cyclonic eddies. Similarly, diverging fronts, indicated by high absolute Lyapunov exponent, appeared to promote the accumulation of larger colonies of Tuff, likely through preferential attachments of individual filaments leading to the generation of long floating rafts at the surface of the Ocean (r=0.33, p-value<0.01).

### Discussion - The spectral biogeography of mesoplankton groups in the Upper Ocean: Implications for present and future ecosystems functioning

In this study, we present a comprehensive analysis linking the size structure of Colonial N_2_-fixers (main autotrophic mesoplankton), Rhizarians (main mixotrophic mesoplankton), and Crustaceans (main heterotrophic mesoplankton), compiled from actual size observations to environmental gradients. We include for the first time diverse iron proxies to model and understand how spectral parameters vary in space and time, alongside traditional drivers like temperature, macronutrient/food availability, or upper ocean mesoscale features. Besides identifying potential drivers of trait distributions in the Global Ocean, our results provide insights into how mesoplankton affect marine ecosystems biogeochemistry and services (N_2_- and CO_2_-fixation, carbon export and micronutrient cycling) in response to environmental stressors. With the predictions of mesoplankton spectral biogeography and associated compilations of pelagic size spectra, expressed as a function of cell abundances or carbon biomass, our hope is to inform the development of biogeochemical and other size-structured models used to foresee the changes in ecosystem functioning and services under future climate change. Since the complexity of plankton functional types represented in the Coupled Model Intercomparison Project 6 models has significantly increased with the support of dataset compilations (e.g MAREDAT, COPEPOD), and plankton size structure, rather than bulk biomass and productivity estimates, is considered the best predictor of ecosystem services (e.g. fish production or carbon export), our dataset should represent a good opportunity to examine the effects of climate change multi-faceted stressors on essential marine ecosystem services.

In essence, our findings revealed that temperature remains a primary factor influencing large-scale shifts in mesoplankton size. Although their overall abundance showed contrasted correlations with sea surface temperature, all groups showed a negative correlation between spectral slopes and sea surface temperature (r(Colonial N_2_-fixers)=-0.23, r(Rhizarians)=-0.29, r(Crustaceans)=-0.48, p-values<0.001), indicating that larger organisms were mostly found in relatively colder environments: a general trend underpinning the Temperature-Size Rule (^23^). The first hypothesis explaining the emergence of larger metazoans in cold environments was linked to increased oxygen availability (^23^), since spatial gradients in surface temperature are opposed to that of dissolved oxygen in marine ecosystems (Supplementary Figure 2). Oxygen exerts a strong metabolic constraint on Crustaceans, with concentrations between 15-50 µM of dissolved oxygen typically defined as critical to maintain aerobic reactions and survival (^42,43^). All Crustacean groups showed a positive correlation between spectral slopes and dissolved oxygen concentration, indicating a flattening of their spectra under high oxygen concentrations. Since endosymbiotic Rhizarians and colonial N_2_-fixers do not require high oxygen concentration to fuel their metabolism (they rather support the production or directly produce oxygen by photosynthesis), it is unlikely that oxygen levels are responsible for the Temperature-Size Rule observed for these groups. Instead, we hypothesize that the apparent Temperature-Size Rule in colonial N_2_-fixers and Rhizarians invokes differences in catabolic (i.e growth) versus anabolic (i.e respiration) rates with respect to changes in temperature. These are typically quantified over a 10°C change, and referred to as temperature coefficients (Q_10_). Anabolic rates should increase more rapidly with temperature compared to catabolic reactions in comparatively smaller organisms, as organisms that present higher Q_10_ coefficients for respiration than for growth (^44^). High respiration rates at high temperatures were shown to support the anoxic conditions that favor N_2_-fixation within colonies of *Trichodesmium* (^45–47)^. Plus, Q_10_ measured from enclosed experiments with Foraminifera (^48^) or Collodaria (^49^) suggest that similar regulatory effects on metabolic reactions may come into play for symbiotic Rhizarians to explain the negative correlation between temperature and their spectral slopes. Our results therefore directly support published meta-analysis (^50^) in suggesting that increased warming and deoxygenation of future marine ecosystems should decrease the size of most mesoplankton groups in the Upper Ocean, likely disrupting both fisheries (^51,52^) and carbon export through reduced gravitational sinking (^53-55)^ and fecal pellet production (^12^).

Our study also highlights several aspects of ecosystem functioning that may limit the negative impacts of global warming and deoxygenation in the Upper Ocean. Compared to previous findings, we captured more variance of mesoplankton trait biogeographies in space and time, with R^2^ of 0.93, 0.84, and 0.66 for N_2_-fixers, Rhizarians and Crustaceans respectively, largely due to the inclusion of various iron proxies underlining the strong linkage between keystone mesoplankton and trace element cycling. Each phylum responded best to a different proxy of iron supply in the epipelagic layer, with a noticeable stimulation of dust and other aeolian iron-rich compounds on the sizes and occurrences of large mixotrophic Rhizarians (e.g., Collodaria, Acantharia) and colonial N_2_-fixers, and a strong link between Crustaceans biogeography and total iron concentrations. While the peak of N_2_-fixing *Trichodesmium* abundance in the Subtropical Atlantic has already been largely attributed to Saharan dust deposition in this ecosystem (^56-58)^, we also found a significant relationship between wet dust deposition and occurrences of Rhizarians . To our knowledge, this relationship has not been demonstrated before and could point to an interesting adaptation of mixotrophic Collodaria and Acantharia, that would require follow-up studies. In the Pacific Ocean, the same groups responded mainly to elevated aerosol fluxes. Phytoplankton blooms, followed by satellite chlorophyll-a concentration, have been regularly linked to direct ash deposition from Southern Pacific tropical islands (Papua New Guinea, Vanuatu and more recently Tonga) volcanic eruptions (^59^), Australian wildfires (^60^), and recently to shallow hydrothermal emissions located across the Tonga-Kermadec volcanic arc (^61^). The latter have been implicated in the iron fertilization of surface waters leading to elevated concentrations of N_2_-fixers (^62^), more specifically *Trichodesmium*, and a shift in the mesoplankton community (^63^). Zooscan analyses highlighted the presence of gelatinous zooplankton in the vicinity of the volcanic arc, in an ecosystem otherwise dominated by copepods. Since nets severely damage the fragile Rhizarians, it was not possible to quantify the impact of iron supply by volcanic emissions, and more generally by aerosols, on this taxonomic group. Our predictions of Rhizarian NBSS appeared linked to aerosol proxy in the SPSG, suggesting that volcanic ashes could also promote the growth of symbiotic Rhizarians like Acantharia and Collodaria. According to the latest CMIP6 simulations, natural dust deposition should increase by 7-20% by the end of the century as a result of enhanced desertification and precipitation as well as changes in wind regimes and land use (^64-66)^, thus potentially promoting carbon and nitrogen fixation in the low and mid latitude ecosystems by mixotrophic Rhizarians and colonial N_2_-fixers. Further, even though aerosol emissions from anthropogenic activities should decline by 50-80% to meet higher air quality standards and the 1.5°C global warming target above pre-industrial level dictated by the Paris agreement by the mid 21^st^ century (^65^), aerosol distribution might also be impacted by alternative natural sources whose emissions could increase up to 125% under global warming according to the CMIP6 models (^66^).

Even though we expected these different iron proxies to be important stressors of mesoplankton biogeography, especially for the autotrophic colonial N_2_-fixers, the ranking of total iron concentration in relation to the spectral biogeography of Crustaceans was the most surprising given that we also included satellite proxies for chlorophyll a concentration or phytoplankton size distribution parameters (slope and fraction of pico-,nano-, and microphytoplankton) in the full model. The coupling between climatologies of satellite microphytoplankton, model total iron concentrations, and predicted Crustacean occurrences in the Equatorial Pacific, California Current upwelling systems, Southern Ocean, subarctic Pacific, and subpolar North Atlantic supports the long-standing recognition that dissolved iron exert a strong control on phytoplankton dynamics (^67^) and associated grazers in the “High Nutrient Low Chlorophyll” (HNLC) ecosystems. Further, the increased concentration of total iron and Crustaceans during and after phytoplankton blooms in polar latitudes highlights the possible role of secondary producers in alleviating nutrient limitation, as noted in Tagliabue et al. (^40^): “higher dissolved iron concentrations associated with high chlorophyll-a could be reconciled by assuming high rates of Fe recycling associated with greater biomass”. Iron fertilization experiments have confirmed the importance of copepods in driving iron recycling and further sustaining phytoplankton blooms in both the Southern (^68^) and the Arctic Ocean (^69^), even though exogenous inputs of iron by river discharges could also be partly responsible for the peak concentration observed after the spring bloom in this ecosystem (^70^). Moreover, a recent comparison between a compilation of total dissolved iron and gridded products (^71^) similar to our analysis has shown that iron was strongly correlated to apparent oxygen utilisation (AOU), which appears positively correlated to Copepod biogeography in our analysis (Supplementary Table 1). These empirical evidence, together with our global analysis linking iron pool to mesozooplankton spectral biogeography, suggest that Crustaceans could maintain elevated iron concentrations by preying on large phytoplankton and remineralizing organic iron into bioavailable forms, further alleviating nutrient limitation of their prey, thus exerting a positive feedback loop that could offset the decline of primary production caused by enhanced stratification and reduced nutrient delivery.

Our comprehensive set of explanatory variables helped build global predictions of mesoplankton spectral biogeography with relatively high R^2^ (0.93, 0.61, and 0.69), providing robust estimates of their size structure related to elemental cycling and ecosystem services and shedding light on novel, important linkages with multiple iron proxies. Future work should focus on addressing the knowledge gaps this study highlights, namely the role of dust and other aerosols in supporting the growth of the mixotrophic Rhizarian classes as well as the role of large Crustaceans in recycling iron in the Upper Ocean, as both processes have the potential to sway the overall decrease of Net Primary Production forecasted in most CMIP6 simulations by the end of the century.

## Material and Methods

### 1. Global compilation of group-specific size spectrum

To understand and predict the global size distribution of Colonial N_2_-fixers, Rhizarians, and Crustaceans in the epipelagic layer, we compiled direct size estimates measured by a range of Underwater Vision Profilers (UVP) of successive generations (^72,73^) and benchtop scanners like the ZooScan (^74^) for nets deployed in all major basins within 0 and 200 m. This compilation stems from a collaborative, international effort, termed the Pelagic Size Structure database (https://pssdb.net), that aims at providing standardized and homogenized size and count information collected by a variety of commercially available plankton imaging instruments. We used the Python workflow (publicly available at https://github.com/jessluo/PSSdb) developed for the first PSSdb release, described in more details in Dugenne et al. (^75^), with little modifications to produce our globally consistent estimates of group-specific Normalized Biovolume Size Spectra (hereafter referred to as NBSS).

Briefly, the general pipeline consists in five automated steps that follow image acquisition and near-real time segmentation (i.e. the process of separating individual particles from background pixels to extract their morphologic features): (1) export of raw size, count and annotation information with the associated metadata from the collaborative platform for image annotation EcoTaxa (https://ecotaxa.obs-vlfr.fr/), (2) standardization of the export tables following a unique format with standard labels and units, (3) quality control of individual datasets that account for particle count uncertainty, sufficient manual validation of the taxonomic classifications (UVP profiles of scanned nets should have ≥95% validated annotations to pass our quality control), differences in size calibration factors, missing information or incorrect GPS location of each sample, (4) aggregation of samples collected in spatial and temporal proximity to provide accurate measures of particle size distribution at the current resolution of biogeochemical models and other global data compilations (i.e. 1°x1° latitude/longitude and year/month), and (5) the computation of NBSS and derived parameters (slope, intercept, R^2^), using pre-defined size classes matching the recent compilation of UVP5 particle size distribution by Kiko et al. (^76^).

The main addition to the Dugenne et al. (^75^) workflow consists of standardizing all taxonomic annotations assigned to individual particles by automated classifiers and later validated by taxonomic experts using the World Register of Marine Species (https://www.marinespecies.org/) as suggested in Neeley et al. (^77^). We wrote a custom script based on existing web scraping functions to automatically search for the taxonomic annotations in WORMS, allowing various formats (e.g. hierarchical or nominal category) and ranks according to the preset classes of automated classifiers, the expertise of human annotators or the research question. The taxonomic look-up table automatically generated by this script is available at https://github.com/jessluo/PSSdb/blob/main/ancillary/plankton_annotated_taxonomy.xlsx. Following this additional standardization, individual annotations were grouped in increasing taxonomic ranks (e.g. phylum, class or subclass, family to assign copepod’s trophic strategy based on published compilation) and according to their morphotypes when possible (e.g. solitary or colonial for Rhizarians, Puff or Tuff for *Trichodesmium)* to estimate group-specific NBSS based on biovolume. Biovolume estimates were computed using equivalent circular diameter (ECD) derived from particles area following equation (1):

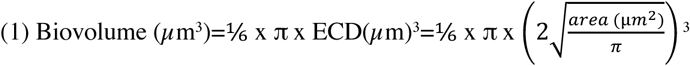

The sum of all biovolume estimates within the discrete size classes defined in equation (2) is then normalized to the volume analyzed, corresponding to the total volume cumulated over depth (0-200 m) and within each spatio-temporal bin, and to the width of each size class to produce NBSS according to equation (3).

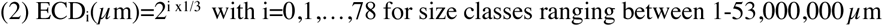

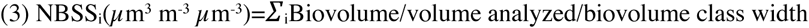

We then selected the maximum and the last normalized biovolume estimate before observing one empty size class respectively as the lower and upper limits of each size spectrum. This step is necessary to select the effective size range of individual mesoplankton group and to ensure that estimates are not biased by the camera resolution or the sampling gear (e.g. net mesh), or by the limited volume sampled respectively. After selection, unbiased NBSS estimates and associated midpoint of individual biovolume size classes are log_10_-transformed to estimate spectral slopes, intercepts, and size ranges for each 1°x1° latitude/longitude and year month by linear regression (Supplementary Fig. 1). Intercept estimates were scaled to the minimum observed size of individual mesoplankton groups since the abundance of 1 µm^3^ particles, the initial intercept for a spectrum starting at log(1 µm^3^)=0, is not representative of the actual abundance of mesoplankton groups. Some of the individual spatio-temporal bins did not contain enough observations to construct a reliable NBSS and fit a log-linear regression as group-specific PSSdb products necessarily included fewer images than the first release. In addition, we selected only datasets that spanned at least 80% of the epipelagic layer (e.g. 160 m) to follow the methodology established by Drago et al. (^78^). As a result, the spatio-temporal coverage in our study is slightly diminished compared to the bulk datasets, constructed with all particles imaged except for methodological artefacts. Overall, group-specific NBSS were computed using 3,068 UVP profiles and 2,411 scans sampled between 2008-2021 and 2004-2022 respectively. These new products can be downloaded at https://doi.org/10.5281/zenodo.10810190.

### 2. Linkages between mesoplankton size structure and environmental covariates

In the absence of broadly concurrent and quality controlled *in situ* hydrographic measurements, we linked mesoplankton NBSS to environmental gradients using global sets of objective analysis, satellite proxies, and biogeochemical models (all accessed and downloaded in Sept. 2023). We used the latest World Ocean Atlas (^79^) objective analysis (WOA, available at https://www.ncei.noaa.gov/access/world-ocean-atlas-2018) to assign each grid cell (defined by 1°x1° latitude/longitude and month) its associated mean mixed layer depth (m), surface temperature (°C), and surface concentration in phosphate (µM), silicate (µM), and nitrate (µM). In addition, we used WOA surface apparent oxygen utilisation (µM) to represent patterns in dissolved oxygen concentration resulting from water mass ventilation and age or remineralization rather than the temperature-dependent O_2_ solubility.

Given that iron is also considered an essential micronutrient for phytoplankton growth (^67^) but *in situ* measurements are still too scarce to construct a reliable distribution of dissolved iron concentrations in the Global Ocean with enough temporal coverage (^80^), we relied on diverse proxies, satellite measurements distributed by NASA; biogeochemical models provided by Copernicus, of both mineral and total iron compounds respectively. In the surface ocean, most labile compounds present are thought to originate from atmospheric aerosols, delivering mineral dust, ashes formed by volcanic activities and wildfires, or anthropogenic compounds all relatively rich in iron (^38^). These can be approximated from space, as total aerosol optical thickness (unitless variable termed Aerosol optical thickness at 869 nm) and wet dust deposition (variable termed dust wet deposition bin 001 or duwt001, in kg m^-2^ s^-1^), with a 9km, monthly resolution using specific satellite algorithms whose products are distributed for every year and month by NASA at https://oceandata.sci.gsfc.nasa.gov/directdataaccess/Level-3%20Mapped/Aqua-MODIS and https://goldsmr4.gesdisc.eosdis.nasa.gov/opendap/MERRA2_MONTHLY/M2TMNXADG.5.12.4/ respectively. Alternatively, organic iron-complexing ligands can also represent important sources of iron, especially for organisms synthesizing siderophore transporters, that are poorly characterized by current analytical methods due to the diversity of compounds and reactivity within this pool (^81^). To account for all compounds in the complex pool of dissolved iron, we used surface predictions of total dissolved iron concentration (µM) from the biogeochemical model PISCES (^82^), distributed for any given year month within the period 1993-2022 by Copernicus (product cmems_mod_glo_bgc_my_0.25_P1M-m available at https://data.marine.copernicus.eu/product/GLOBAL_MULTIYEAR_BGC_001_029/) with a resolution of 1/4° latitude/longitude. This model has shown a strong agreement with benchmark sections from the GEOTRACES program compared to other model predictions and also include all known source and sink processes (riverine, sediment, hydrothermal, or dust inputs, remineralization, scavenging) affecting iron pools in marine ecosystems (^80^).

We downloaded monthly average surface photosynthetically available radiation (PAR, in E m^-2^ d^-1^) and chlorophyll-a concentration (CHL, in mg m^-3^) for all sampled years (2004-2022) from the same url as the aerosol NASA satellite products. Besides chlorophyll-a concentration, we also included recent estimates of the particle size distribution slope (PSD slope, in particles L^-1^ µm^-2^) and fractions of pico-(0.2-2 µm), nano-(2-20 µm), and micro-(20-50 µm) phytoplankton retrieved from backscatter coefficients measured by remote sensing at a resolution 4km for every month year (^83^) to characterize the phytoplankton community representing the main food source for mesozooplankton groups. Lastly, proxies for mesoscale structure, like sea level anomaly (SLA, in m), that indicate cyclonic and anticyclonic eddies under negative and positive anomaly, and finite size Lyapunov exponent (FSLE, in d^-1^), that measures the frontal divergence of mesoscale eddies or other currents in close proximity, were accessed on AVISO at https://www.aviso.altimetry.fr/en/data/products with a resolution of 1/25° latitude/longitude. Since mesoscale features are operationally defined as coherent vortex generally trackable for a month or longer with a radius scale of ∼100 km (^84^), we consider that the final 1°x1° (∼111km) and year month resolution of our datasets were also representative of mesoscale circulation, even at the higher end member of its operational resolution.

All variables were automatically downloaded and averaged to match the resolution of our mesoplankton NBSS estimates (if the resolution was greater than 1°x1° latitude/longitude and year month) using custom scripts available at https://github.com/mdugenne/PSSdb_Learning. Resulting spatial patterns are presented in Supplementary Fig. 2. The environmental datasets were then merged to our group-specific products according to the bin location (1°x1° latitude/longitude) and the sampling time (year month for yearly resolved variables or month for WOA climatologies). Locations with one or more missing environmental variable(s) (e.g. satellite products at high latitude during winter) were filtered out to compute the Spearman correlation coefficients between environmental factors and spectral parameters (slope, intercept, size range) and train the machine learning model used to predict the size distribution of mesoplankton groups globally.

### 3. Global predictions of mesoplankton size structure using machine learning models

We used our compilations of group-specific NBSS and associated environmental factors in combination with machine learning habitat models to estimate the global size distribution of the three major mesoplankton groups (Colonial N_2_-fixers, Rhizarians, Crustaceans) in the epipelagic layer. Like Barton et al. (^21^), we recognize that mesoplankton size biogeography is ultimately determined by the interplay between both bottom-up and top-down (e.g. size-selective predation by fish or other nekton) controls, though the latter are out of this paper’s scope as this would require an extensive characterization of larger predators (through space and time) that are poorly detected with the imaging systems we used.

Our modelling effort builds on the recent publication of Drago et al. (^78^), who characterized and predicted the total biomass estimates of several mesozooplankton groups (e.g. Copepods, Rhizarians, not including colonial N_2_-fixers) using UVP profiles globally distributed. Since elevated biomass may result from increased abundance and/or size, we expanded their work to help disentangle the effect of environmental factors on mesoplankton abundance (with the maximum abundance representing the spectral intercept) and size (whose continuum is determined by spectral slopes and effective range). Habitat models cover diverse statistical models, including the relatively recent boosted regression trees (also referred as xgboost,^85^), that link environmental factors to species or group distribution with the objective to predict their occurrence in unsampled locations. Like other machine learning models, xgboost learns simple or complex rules (here a hierarchy of yes/no decisions delimited by the maximum tree depth) linking the response variable (NBSS parameters) to explanatory variables (environmental factors described in Material and Methods section 2) using a subset of the dataset (here 80% random split), named the training set. The learning process is an iterative process based on the progressive evaluation of a loss function, defined as the root mean square error between observations and predictions.

Their performance is then assessed using the remaining subset (here 20% random split), termed the test set. The algorithm performance evaluation is based on different metrics like the coefficient of determination (R^2^), determined by the sum of squared errors between test predictions and test set, or the log-likelihood of the model, that represents the goodness of fit between model predictions and observations according to the data distribution. We used two parameters linked to the model log-likelihood (noted *l*), the Akaike (AIC) and Bayesian (BIC) Information Criterion defined in equation (4), to compare model predictions amongst increasing taxonomic ranks. While we expect that grouping observations based on high taxonomic rank like class or families should be more appropriate to capture the variability in mesoplankton NBSS on a global scale as a result of diverging environmental adaptations (^24^), we also considered that the decrease of available observations for these ranks or the differences in model complexity could also result in overall lower performances of the model. Since AIC and BIC take into account model likelihood, complexity, and the number of available observations, we used these estimates as a mean to provide objective comparisons between model performances. The likelihood of taxon-specific models were averaged, with a weight corresponding to the proportion of observations, across all subgroups of colonial N2-fixers, Rhizarians and Crustaceans when increased taxonomic ranks were compared to the overall model performed at the phylum level, following a similar approach as mixture models (^86^).

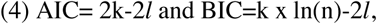

with k the number of model parameters, n the number of observations, and *l* the model log-likelihood directly returned by the xgboost package.

Unlike more traditional decision tree models that predict a set of discrete values termed the leaves, xgboost models may apply to continuous variables by producing numerous leaves associated with a specific score, that are ultimately summed up across all trees to create a smooth gradient in the predicted response variable (i.e the “boosting” process). Instead of discontinuous response, the resulting predictions will thus present a smoother data distribution similar to that of spectral parameters (Supplementary Fig. 1).

Since these models are based on direct representation of the linkages between explanatory and response variables, estimates of feature importance, measured as the average gain in model performance (log-likelihood) across all splits after the variable was used, are relatively straightforward. Such models have additional advantages as they can deal with many dependent variables (for example, dust and aerosol proxies for iron deposition, or temperature and PAR) and predict multivariate outputs. In our study, we simultaneously predicted spectral slopes, intercepts and size range while accounting for their correlation (Supplementary Fig. 1).

The reconstructed NBSS, obtained by assuming a log-linear decline with the modeled parameters, can then be integrated to predict the total abundances of mesoplankton groups at any given space and time. Prior to model training, both the spectral intercept and size range were log_10_-transformed to ensure that predictions will be positive after reverse transformation. To minimize the effect of irrelevant explanatory variables and avoid model overfitting, several parameters were tuned, including score shrinkage (eta parameter tested between 0.1 and 0.5) to homogenize the gradient boosting process in case of overfitting, the maximum number of splits in the decision trees (max_depth parameter tested between 3 and 10) that can lead to oversimplification/complexification of the model, or the number of trees (n_estimators parameter tested between 20 and 100). The best combination of model parameters was determined objectively using the grid search function implemented in the xgboost package.

Not all mesoplankton groups were abundant enough or significantly correlated with the environmental factors tested to analyze and predict their size structure in the epipelagic layer using a machine learning model (^78^). Overall, we could accurately predict (i.e., with high R^2^) the spectral biogeography of one family of colonial N_2_-fixers (Oscillatoriales), four classes of Rhizarians (Acantharia, Collodaria, Phaeodaria, Foraminifera) and three classes of Crustaceans (Copepoda, Malacostraca, Ostracoda). The family of colonial N_2_-fixers was represented exclusively by *Trichodesmium* in this study although we note the presence of a few Nostocales in the Baltic Sea that were too rare for reliable predictions of their size spectrum (data not shown). Some groups were further distinguished based on their morphology (e.g. *Trichodesmium* puff or tuff), or into functional groups (e.g. carnivorous or omnivorous copepods) based on published correspondence between copepod’s families and trophic groups (^41^). The most abundant class of Rhizaria, Collodaria, were almost exclusively found as solitary forms in the epipelagic, so we could not explore possible niche partitioning amongst colonial and solitary Rhizarians. Predictions were generated with the optimal set of parameters in every 1°x1° grid cell for the full temporal coverage of the UVP and scanners datasets. Temporal correlations with the multiple iron proxies (total iron, dust or aerosol) were estimated in each spatial bin to determine which proxy had the strongest link with NBSS integral predictions, that result from predicted slopes, intercepts and size ranges (Supplementary Fig. 3). The main iron proxy was then defined based on the maximum correlation across the 3 possible correlation coefficients in broad ecosystem categories (e.g. Polar, Temperate-Subpolar, Subtropical Gyres, Equatorial-Upwellings) corresponding to adjacent Longhurst provinces (https://www.marineregions.org/sources.php#longhurst), and in diverse ocean basins (https://www.marineregions.org/sources.php#goas). For a full description of the xgboost python package used in this study, see documentation provided at https://xgboost.readthedocs.io/en/stable/python. The code to train, check the model performance and feature importance, and predict mesoplankton size distribution based on xgboost functions is available at https://github.com/mdugenne/PSSdb_Learning.

## Acknowledgments

This work was primarily funded by NOAA, award NA21OAR4310254 (JL, RK, LS, FL, J-OI, TO’B and CS), in addition to the following funding programs: European Union Horizon 2020 program grant 817578 (MD, RK, and LS)

French National Research Agency (ANR), Programme d’Investissements d’Avenir grant #ANR-19-MPGA-0012 (RK) Heisenberg Programme of the German Science Foundation #KI1387/5-1 (RK).

## Author contributions

Conceptualization: MD, MC-U, JYL, RK, TO’B, J-OI, FL, LS, CS Methodology: MD, MC-U, JYL, RK Data acquisition: RK, J-OI, FL, LS, AC, LG, CG, HH, AMcD, MN, MP, and J-BR

Data collection, acquisition, or curation: NB, SB, FC, CD, LD, AE, NG, JAH, LJ, ML, ZM, BN, TP, ER, JT, CT, MV Visualization: MD Supervision: JYL, RK Writing—original draft: MD. Writing—review & editing: all authors

Competing interests: Authors declare that they have no competing interests

PSSdb data contributors consortium: Marc Picheral (Sorbonne Université, CNRS, Laboratoire d’Océanographie de Villefranche, Villefranche-sur-mer, France), Lionel Guidi (Sorbonne Université, CNRS, Laboratoire d’Océanographie de Villefranche, Villefranche-sur-mer, France), Andrew M. P. McDonnell (Oceanography Department, University of Alaska Fairbanks, Fairbanks, AK, United States), Laetitia Drago (Sorbonne Université, CNRS, Laboratoire d’Océanographie de Villefranche, Villefranche-sur-mer, France), Zoé Mériguet (Sorbonne Université, CNRS, Laboratoire d’Océanographie de Villefranche, Villefranche-sur-mer, France), Marion Vilain (Sorbonne Université, CNRS, Laboratoire d’Océanographie de Villefranche, Villefranche-sur-mer, France), Chloé Tilliette (Sorbonne Université, CNRS, Laboratoire d’Océanographie de Villefranche, Villefranche-sur-mer, France), Nagib Bhairy (Mediterranean Institute of Oceanography, Marseille, France), Sophie Bonnet (Mediterranean Institute of Oceanography, Marseille, France), Thelma Panaïotis (Sorbonne Université, CNRS, Laboratoire d’Océanographie de Villefranche, Villefranche-sur-mer, France), Margaux Noyon (Department of Oceanography and Institute for Coastal and Marine Research, Nelson Mandela University, Gqeberha, South Africa), Cécile Guieu (Sorbonne Université, CNRS, Laboratoire d’Océanographie de Villefranche, Villefranche-sur-mer, France), Astrid Cornils (Polar Biological Oceanography, Alfred Wegener Institute Helmholtz Centre for Polar and Marine Research, Bremerhaven, Germany), Jean-Baptiste Romagnan (DECOD, IFREMER, INRAE, Institut Agro, Nantes, France), Nina Grandrémy (DECOD, IFREMER, INRAE, Institut Agro, Nantes, France), François Carlotti (Mediterranean Institute of Oceanography, Marseille, France), Magali Lescot (Mediterranean Institute of Oceanography, Marseille, France), Emilie Riquier (Sorbonne Université, CNRS, Laboratoire d’Océanographie de Villefranche, Villefranche-sur-mer, France), Laetitia Jalabert (Sorbonne Université, CNRS, Laboratoire d’Océanographie de Villefranche, Villefranche-sur-mer, France), Amanda Elineau (Sorbonne Université, CNRS, Laboratoire d’Océanographie de Villefranche, Villefranche-sur-mer, France), Corinne Desnos (Sorbonne Université, CNRS, Laboratoire d’Océanographie de Villefranche, Villefranche1280 sur-mer, France), Helena Hauss (GEOMAR Helmholtz Centre for Ocean Research Kiel, Kiel, Germany), Jan Taucher (GEOMAR Helmholtz Centre for Ocean Research Kiel, Kiel, Germany), Barbara Niehoff (Polar Biological Oceanography, Alfred Wegener Institute Helmholtz Centre for Polar and Marine Research, Bremerhaven, Germany), Jenny Huggett (Oceans and Coastal Research, Department of Forestry, Fisheries and the Environment, Cape Town, South Africa)

## Supplementary information

**Supplementary Figure 1.**
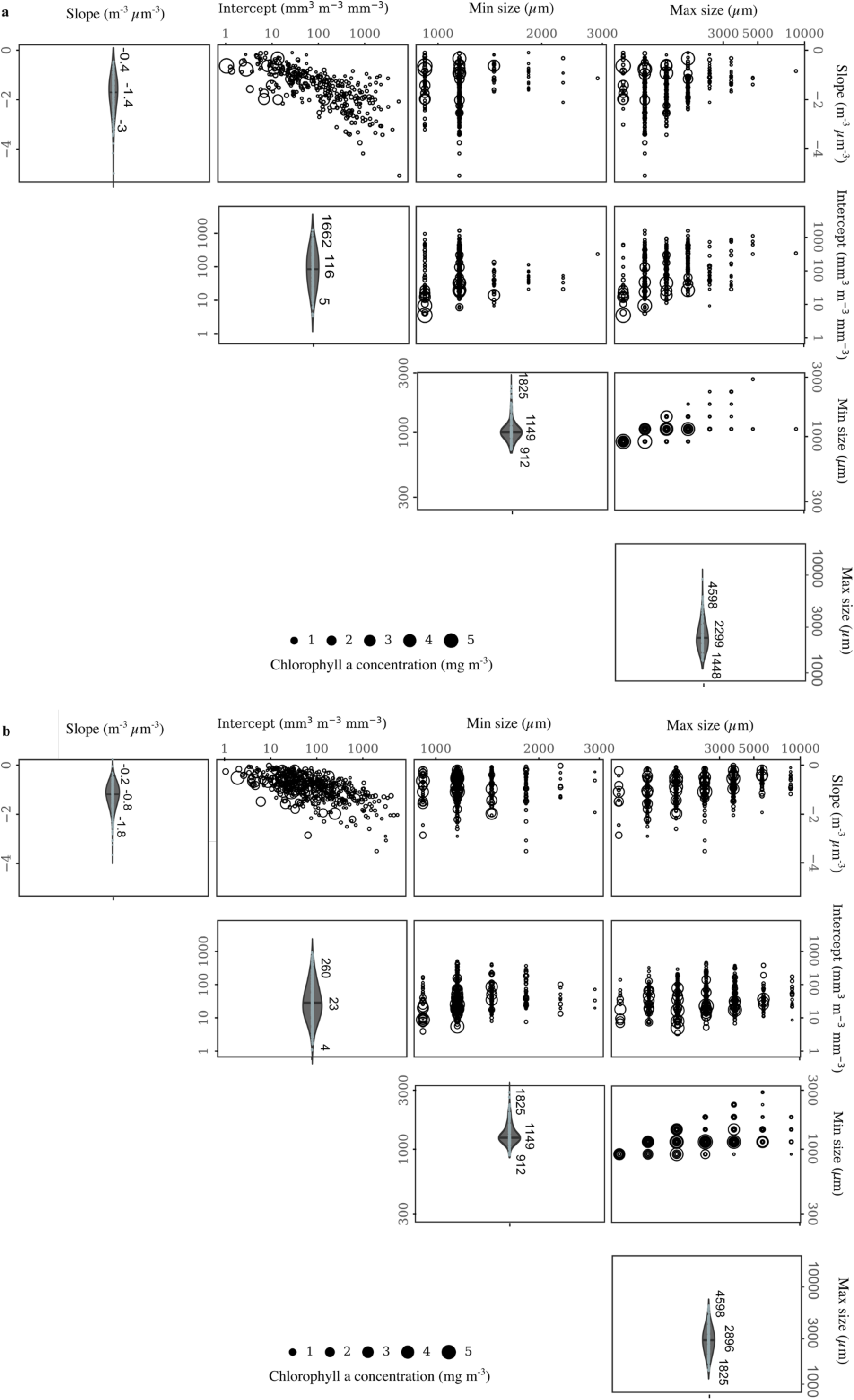
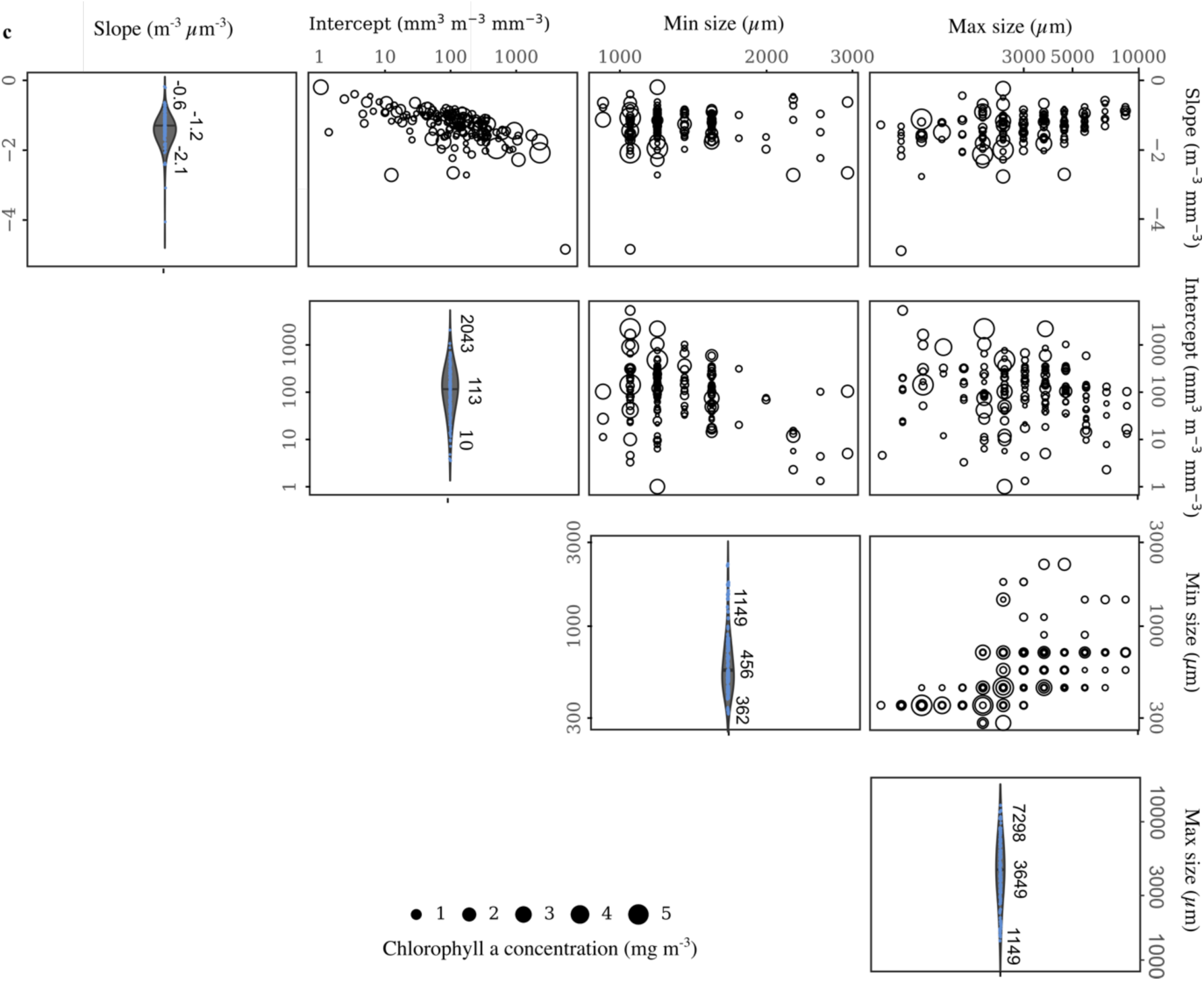
Distribution (diagonal) and correlation within spectral parameters (intercept, slope, and size range) of three mesoplankton groups. a Colonial N_2_-fixers. b Rhizarians. c Crustaceans. Datapoints vary in size according to the average chlorophyll-a concentration observed in satellite products in the same spatio-temporal bin. The distribution of spectral parameters is shown with violin plots on the diagonal.

**Supplementary Figure 2.**
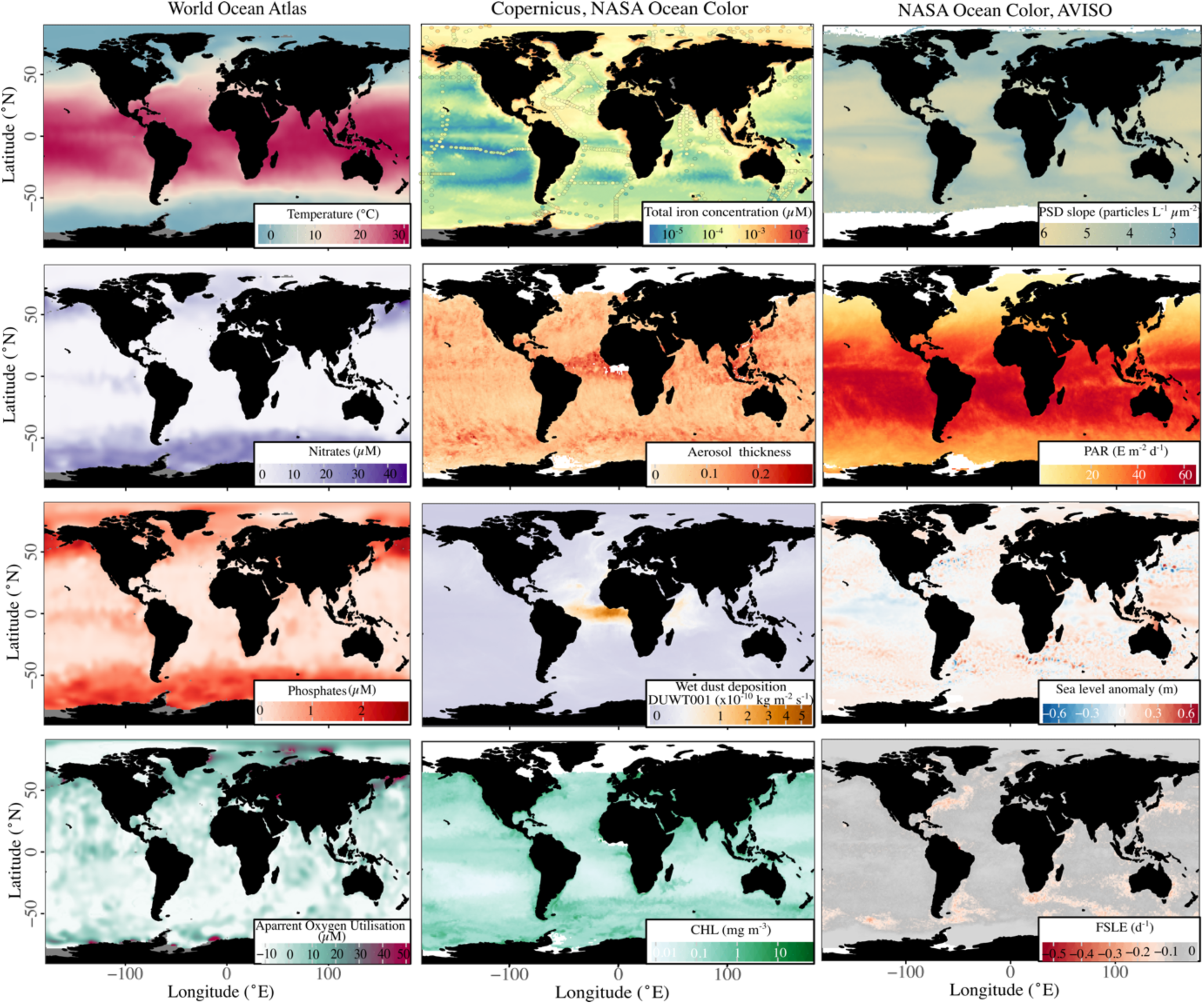
Environmental factors used to predict the spectral biogeography of the main mesoplankton groups. Set of average (for the month of January) environmental factors included as explanatory variables to predict the size distribution of colonial N_2_-fixers, Rhizarians and Crustaceans globally. Variables include surface temperature and concentrations of nitrates, phosphates, silicates (not shown given its strong correlation with nitrates), and apparent oxygen utilization, along with mixed layer depth (MLD) from World Ocean Atlas. The surface concentration of diverse iron pools were approximated by satellite (wet dust deposition and aerosol optical thickness) or biogeochemical model (total iron) products. Total iron concentrations measured *in situ* as part of the GEOTRACES program (colored dots, downloaded at https://geotraces.webodv.awi.de/service/) are superimposed on top of the model PISCES continuous predictions (distributed by Copernicus), to validate the product used in our study. Phytoplankton concentration and composition was approximated by chlorophyll-a concentration (CHL), particle size distribution slope (PSD slope), and fractions of pico-, nano-and microphytoplankton (not shown since their relative proportion affect directly PSD slopes) satellite products distributed by NASA, alongside daily photosynthetically available radiance (PAR). Mesoscale activities were indicated by two AVISO products: Sea level anomaly (SLA), indicative of cyclonic or anticyclonic eddies under negative or positive anomaly, and finite size Lyapunov exponent (FSLE), marking the divergence rate of water parcels in close proximity.

**Supplementary Figure 3.**
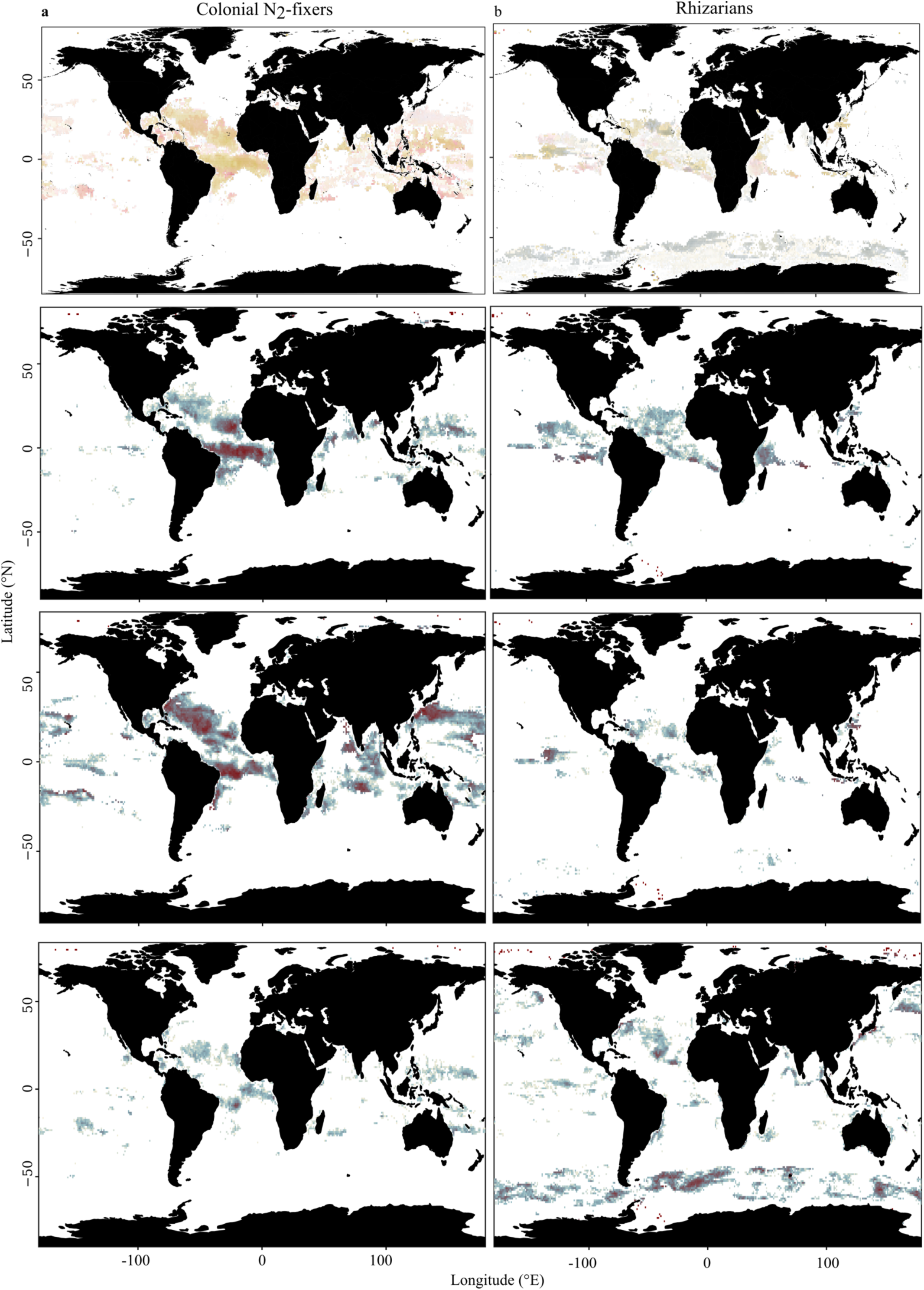
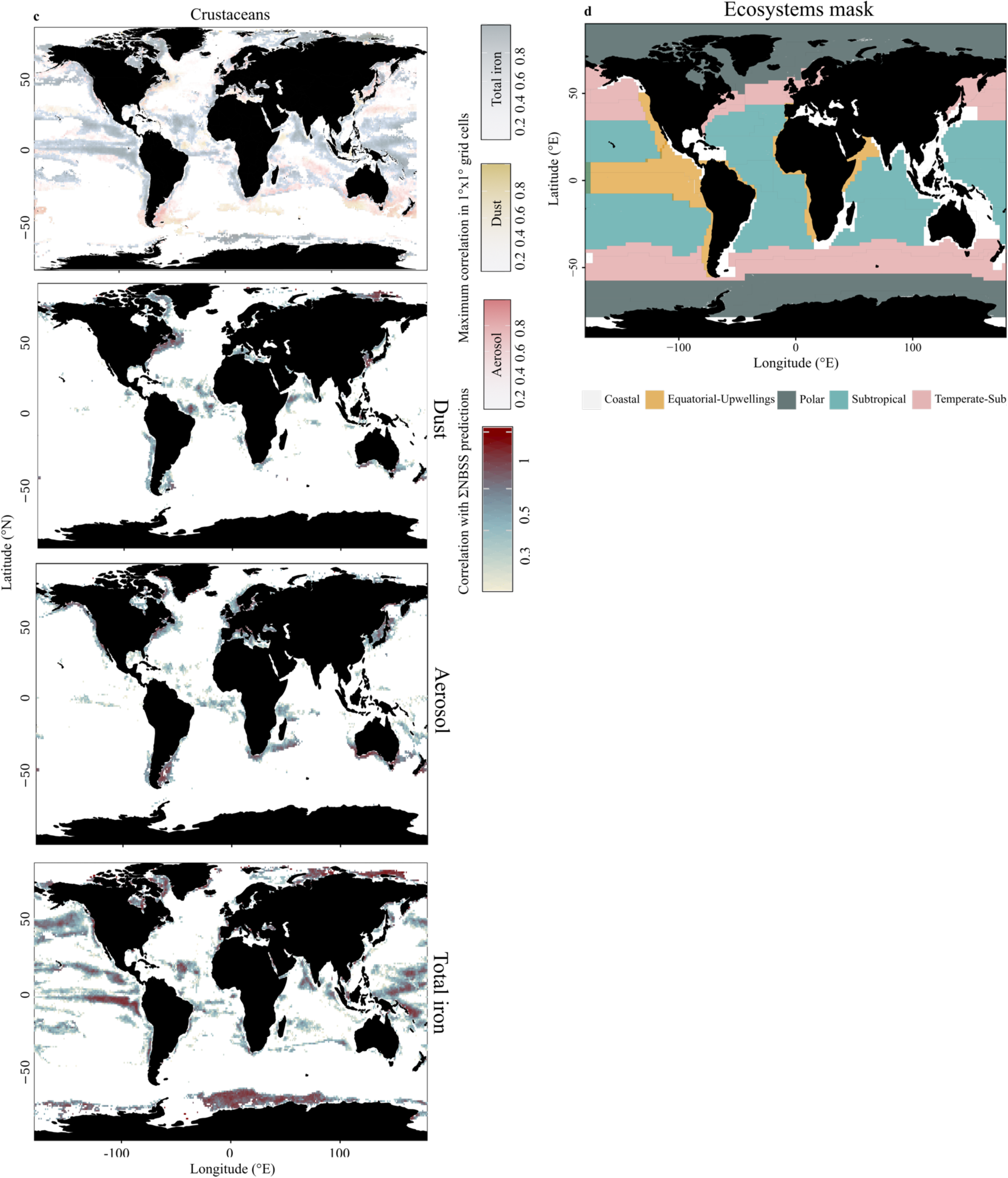
Correlations between iron proxies and predictions of NBSS integral in 1°x1° grid cells. **a** Identification of main iron proxy (top panel) based on the maximum correlation coefficients between iron proxies (dust, aerosol, or total iron) and predictions of colonial N2-fixers NBSS integral in 1°x1° grid cells. **b** Identification of main iron proxy (top panel) based on the maximum correlation coefficients between iron proxies (dust, aerosol, or total iron) and predictions of Rhizarians NBSS integral in 1°x1° grid cells. **c** Identification of main iron proxy (top panel) based on the maximum correlation coefficients between iron proxies (dust, aerosol, or total iron) and predictions of Crustaceans NBSS integral in 1°x1° grid cells. **d** Mask of ecosystems (Equatorial-Upwellings: orange, Polar: green, Subtropical: blue, Temperate-Subpolar: pink) used to plot seasonal trends of NBSS predictions in association with the main iron proxy. The main proxy is identified in each ecosystem / ocean basins by calculating the number of bins associated to each iron proxy. The proxy with the maximum number of bins in a given ecosystem/ocean basin is considered d the main iron pool.

**Supplementary Table 1.**
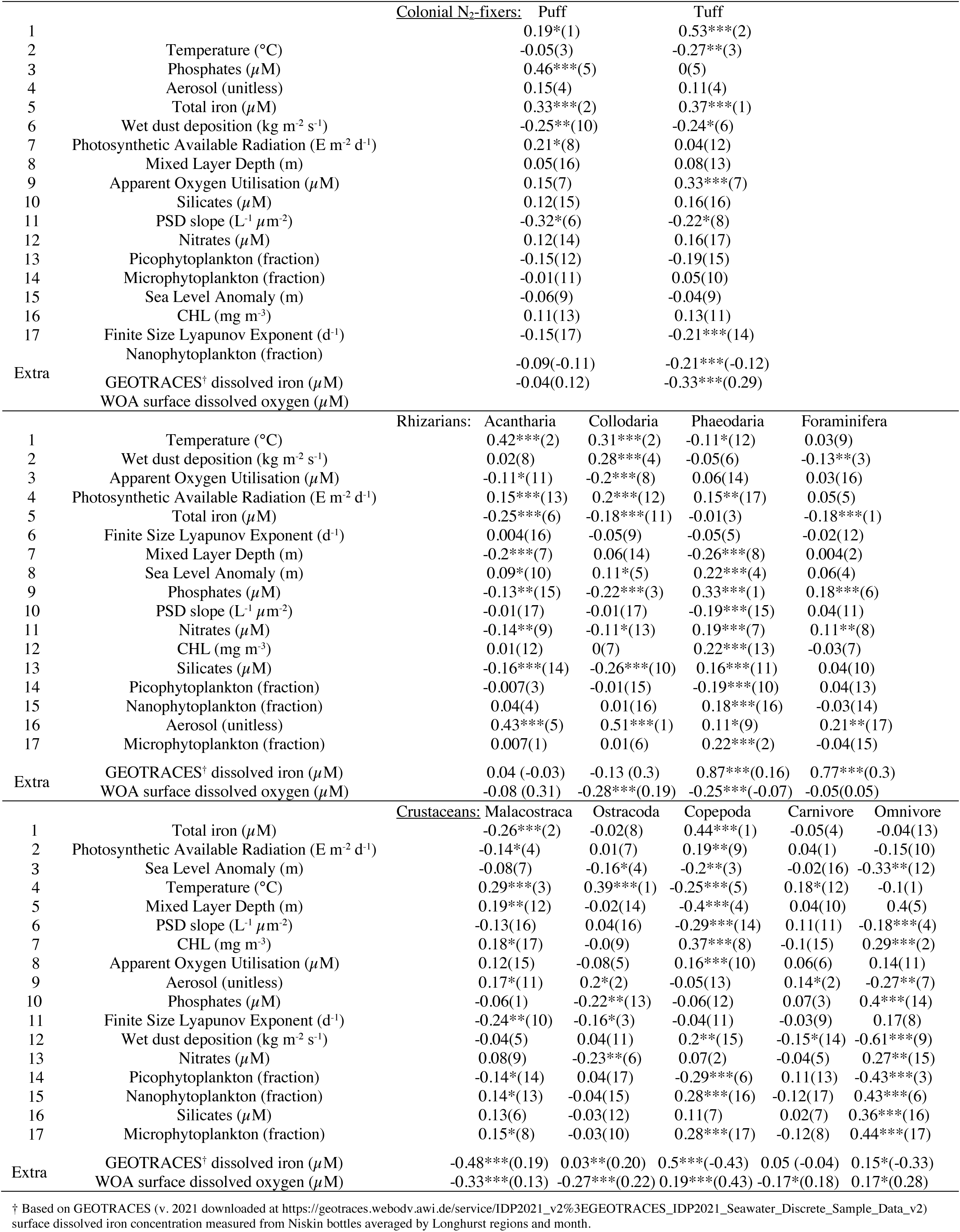
Spearman correlation coefficients between environmental factors and the integral of mesoplankton size spectra culated at the finest taxonomic resolution. Environmental factors are ranked according to their feature importance in model fits.

## References

1. Sheldon, R. W., Prakash, A. & Sutcliffe Jr., W. H. The size distribution of particles in the Ocean. Limnology and Oceanography 17, 327–340 (1972).

2. Platt, T. & Denman, K. Organisation in the pelagic ecosystem. Helgolander Wiss. Meeresunters 30, 575–581 (1977).

3. Petchey, O. L. & Belgrano, A. Body-size distributions and size-spectra: universal indicators of ecological status? Biology Letters 6, 434–437 (2010).

4. Trebilco, R., Baum, J. K., Salomon, A. K. & Dulvy, N. K. Ecosystem ecology: size-based constraints on the pyramids of life. 21Trends in Ecology & Evolution 28, 423–431 (2013).

5. Henson, S., Le Moigne, F. & Giering, S. Drivers of Carbon Export Efficiency in the Global Ocean. Global Biogeochemical Cycles 33, 891–903 (2019).

6. Serra-Pompei, C. et al. Linking Plankton Size Spectra and Community Composition to Carbon Export and Its Efficiency. Global Biogeochemical Cycles 36, e2021GB007275 (2022).

7. Sheldon, R. W., Sutcliffe Jr., W. H. & Paranjape, M. A. Structure of Pelagic Food Chain and Relationship Between Plankton and Fish Production. J. Fish. Res. Bd. Can. 34, 2344–2353 (1977).

8. Rykaczewski, R. R. & Checkley, D. M. Influence of ocean winds on the pelagic ecosystem in upwelling regions. Proceedings of the National Academy of Sciences 105, 1965–1970 (2008).

9. Maranón, E., Cermeóo, P., Rodríguez, J., Zubkov, M. V. & Harris, R. P. Scaling of phytoplankton photosynthesis and cell size in the ocean. Limnology and Oceanography 52, 2190–2198 (2007).

10. Kiørboe, T. & Hirst, A. G. Shifts in Mass Scaling of Respiration, Feeding, and Growth Rates across Life-Form Transitions in Marine Pelagic Organisms. The American Naturalist 183, E118–E130 (2014).

11. Hansen, B., Bjornsen, P. K. & Hansen, P. J. The size ratio between planktonic predators and their prey. Limnology and Oceanography 39, 395–403 (1994).

12. Stamieszkin, K. et al. Size as the master trait in modeled copepod fecal pellet carbon flux. Limnology and Oceanography 60, 2090–2107 (2015).

13. Uye, S. & Kaname, K. Relations between fecal pellet volume and body size for major zooplankters of the Inland Sea of Japan. J Oceanogr 50, 43–49 (1994).

14. Chindia, J. A. & Figueredo, C. C. Phytoplankton settling depends on cell morphological traits, but what is the best predictor? Hydrobiologia 813, 51–61 (2018).

15. Andersen, K. H. et al. Characteristic Sizes of Life in the Oceans, from Bacteria to Whales. Annual Review of Marine Science 8, 217–241 (2016).

16. Blanchard, J. L., Heneghan, R. F., Everett, J. D., Trebilco, R. & Richardson, A. J. From Bacteria to Whales: Using Functional Size Spectra to Model Marine Ecosystems. Trends in Ecology & Evolution 32, 174–186 (2017).

17. Hatton, I. A., Heneghan, R. F., Bar-On, Y. M. & Galbraith, E. D. The global ocean size spectrum from bacteria to whales. Science Advances 7, eabh3732 (2021).

18. Heneghan, R. F., Hatton, I. A. & Galbraith, E. D. Climate change impacts on marine ecosystems through the lens of the size spectrum. Emerging Topics in Life Sciences 3, 233–243 (2019).

19. Pierella Karlusich, J. J., et al. Global distribution patterns of marine nitrogen-fixers by imaging and molecular methods. Nat Commun 12, 4160 (2021).

20. Brandão, M. C. et al. Macroscale patterns of oceanic zooplankton composition and size structure. Sci Rep 11, 15714 (2021).

21. Barton, A. D. et al. The biogeography of marine plankton traits. Ecology Letters 16, 522–534 (2013).

22. Panaïotis, T. et al. Three major mesoplanktonic communities resolved by in situ imaging in the upper 500 m of the global ocean. Global Ecology and Biogeography **n/a**,.

23. Atkinson, D. & Sibly, R. M. Why are organisms usually bigger in colder environments? Making sense of a life history puzzle. Trends in Ecology & Evolution 12, 235–239 (1997).

24. Brun, P., Payne, M. R. & Kiørboe, T. Trait biogeography of marine copepods – an analysis across scales. Ecology Letters 19, 1403–1413 (2016).

25. Wrightson, L. & Tagliabue, A. Quantifying the Impact of Climate Change on Marine Diazotrophy: Insights From Earth System Models. Front. Mar. Sci. 7, (2020).

26. Faure, E. et al. Mixotrophic protists display contrasted biogeographies in the global ocean. ISME J 13, 1072–1083 (2019).

27. Laget, M. et al. Global census of the significance of giant mesopelagic protists to the marine carbon and silicon cycles. Nat Commun 15, 3341 (2024).

28. Beck, M. et al. Morphological diversity increases with decreasing resources along a zooplankton time series. Proceedings of the Royal Society B: Biological Sciences 290, 20232109 (2023).

29. Vilgrain, L. et al. Trait-based approach using in situ copepod images reveals contrasting ecological patterns across an Arctic ice melt zone. Limnology and Oceanography 66, 1155–1167 (2021).

30. Kiørboe, T., Visser, A. & Andersen, K. H. A trait-based approach to ocean ecology. ICES Journal of Marine Science 75, 1849–1863 (2018).

31. Litchman, E., Ohman, M. D. & Kiørboe, T. Trait-based approaches to zooplankton communities. Journal of Plankton Research 35, 473–484 (2013).

32. Ohman, M. D. & Romagnan, J.-B. Nonlinear effects of body size and optical attenuation on Diel Vertical Migration by zooplankton. Limnology and Oceanography 61, 765–770 (2016).

33. Litchman, E., Klausmeier, C. A., Schofield, O. M. & Falkowski, P. G. The role of functional traits and trade-offs in structuring phytoplankton communities: scaling from cellular to ecosystem level. Ecology Letters 10, 1170–1181 (2007).

34. Martini, S. et al. Functional trait-based approaches as a common framework for aquatic ecologists. Limnology and Oceanography 66, 965–994 (2021).

35. Orenstein, E. C. et al. Machine learning techniques to characterize functional traits of plankton from image data. Limnology and Oceanography 67, 1647–1669 (2022).

36. Lombard, F. et al. Globally consistent quantitative observations of planktonic ecosystems. Frontiers in Marine Science 6, (2019).

37. Kiko, R., Lopes, R. M., Soviadan, Y. D. & Stemmann, L. Towards a distributed and operational pelagic imaging network. Ocean Coast. Res. 71, e23058 (2023).

38. Tagliabue, A. et al. The integral role of iron in ocean biogeochemistry. Nature 543, 51–59 (2017).

39. Wang, L.-C. & Jia-Yuh, Y. Dynamics of Upwelling Annual Cycle in the Equatorial Pacific Ocean. Sustainability 11, 5038 (2019).

40. Tagliabue, A. et al. A global compilation of dissolved iron measurements: focus on distributions and processes in the Southern Ocean. Biogeosciences 9, 2333–2349 (2012).

41. Benedetti, F., Wydler, J. & Vogt, M. Copepod functional traits and groups show divergent biogeographies in the global ocean. Journal of Biogeography 50, 8–22 (2023).

42. Seibel, B. A., Schneider, J. L., Kaartvedt, S., Wishner, K. F. & Daly, K. L. Hypoxia Tolerance and Metabolic Suppression in Oxygen Minimum Zone Euphausiids: Implications for Ocean Deoxygenation and Biogeochemical Cycles. Integrative and Comparative Biology 56, 510–523 (2016).

43. Kiko, R., Hauss, H., Buchholz, F. & Melzner, F. Ammonium excretion and oxygen respiration of tropical copepods and euphausiids exposed to oxygen minimum zone conditions. Biogeosciences 13, 2241–2255 (2016).

44. Regaudie-de-Gioux, A. & Duarte, C. M. Temperature dependence of planktonic metabolism in the ocean. Global Biogeochemical Cycles 26, (2012).

45. Staal, M., Meysman, F. J. R. & Stal, L. J. Temperature excludes N2-fixing heterocystous cyanobacteria in the tropical oceans. Nature 425, 504–507 (2003).

46. Stal, L. J. Is the distribution of nitrogen-fixing cyanobacteria in the oceans related to temperature? Environmental Microbiology 11, 1632–1645 (2009).

47. Inomura, K., Wilson, S. T. & Deutsch, C. Mechanistic Model for the Coexistence of Nitrogen Fixation and Photosynthesis in Marine Trichodesmium. mSystems 4, e00210–19 (2019).

48. Lombard, F., Erez, J., Michel, E. & Labeyrie, L. Temperature effect on respiration and photosynthesis of the symbiont-bearing planktonic foraminifera Globigerinoides ruber, Orbulina universa, and Globigerinella siphonifera. Limnology and Oceanography 54, 210–218 (2009).

49. Villar, E. et al. Symbiont Chloroplasts Remain Active During Bleaching-Like Response Induced by Thermal Stress in Collozoum pelagicum (Collodaria, Retaria). Frontiers in Marine Science 5, (2018).

50. Daufresne, M., Lengfellner, K. & Sommer, U. Global warming benefits the small in aquatic ecosystems. Proc Natl Acad Sci U S A 106, 12788–12793 (2009).

51. Rice, E., Dam, H. G. & Stewart, G. Impact of Climate Change on Estuarine Zooplankton: Surface Water Warming in Long Island Sound Is Associated with Changes in Copepod Size and Community Structure. Estuaries and Coasts 38, 13–23 (2015).

52. Heneghan, R. F., Everett, J. D., Blanchard, J. L., Sykes, P. & Richardson, A. J. Climate-driven zooplankton shifts cause large-scale declines in food quality for fish. Nat. Clim. Chang. 13, 470–477 (2023).

53. Guidi, L. et al. Plankton networks driving carbon export in the oligotrophic ocean. Nature 532, 465–470 (2016).

54. Agusti, S. et al. Ubiquitous healthy diatoms in the deep sea confirm deep carbon injection by the biological pump. Nat Commun 6, 7608 (2015).

55. Benavides, M. et al. Sinking Trichodesmium fixes nitrogen in the dark ocean. The ISME Journal 16, 2398–2405 (2022).

56. Tovar-Sanchez, A. et al. Effects of Dust Deposition and River Discharges on Trace Metal Composition of Trichodesmium spp. in the Tropical and Subtropical North Atlantic Ocean. Limnology and Oceanography 51, 1755–1761 (2006).

57. Fernández, A., Mouriño-Carballido, B., Bode, A., Varela, M. & Marañón, E. Latitudinal distribution of *Trichodesmium* spp. and N_2_ fixation in the Atlantic Ocean. Biogeosciences 7, 3167–3176 (2010).

58. Langlois, R. J., Mills, M. M., Ridame, C., Croot, P. & LaRoche, J. Diazotrophic bacteria respond to Saharan dust additions. Marine Ecology Progress Series 470, 1–14 (2012).

59. Yoon, J.-E., King, D., Longman, J. & Cronin, S. J. Differential response of chlorophyll-a concentrations to explosive volcanism in the western South Pacific. Frontiers in Marine Science 10, (2023).

60. Li, M., Shen, F. & Sun, X. 2019 ‒2020 Australian bushfire air particulate pollution and impact on the South Pacific Ocean. Sci Rep 11, 12288 (2021).

61. Guieu, C. et al. Iron from a submarine source impacts the productive layer of the Western Tropical South Pacific (WTSP). Sci Rep 8, 9075 (2018).

62. Bonnet, S. et al. Natural iron fertilization by shallow hydrothermal sources fuels diazotroph blooms in the ocean. Science 380, 812–817 (2023).

63. Mériguet, Z. et al. Plankton community structure in response to hydrothermal iron inputs along the Tonga-Kermadec arc. Front. Mar. Sci. 10, 1232923 (2023).

64. Liu, J. et al. Historical footprints and future projections of global dust burden from bias-corrected CMIP6 models. *npj Clim Atmos Sci* **7**, 1–12 (2024).

65. Wang, P. et al. Aerosols overtake greenhouse gases causing a warmer climate and more weather extremes toward carbon neutrality. Nat Commun 14, 7257 (2023).

66. Gomez, J. et al. The projected future degradation in air quality is caused by more abundant natural aerosols in a warmer world. Commun Earth Environ 4, 1–11 (2023).

67. Moore, C. M. et al. Processes and patterns of oceanic nutrient limitation. Nature Geosci 6, 701–710 (2013).

68. Laglera, L. M. et al. Iron partitioning during LOHAFEX: Copepod grazing as a major driver for iron recycling in the Southern Ocean. Marine Chemistry 196, 148–161 (2017).

69. Tsuda, A. et al. Evidence for the grazing hypothesis: Grazing reduces phytoplankton responses of the HNLC ecosystem to iron enrichment in the western subarctic pacific (SEEDS II). J Oceanogr 63, 983–994 (2007).

70. Winkelbauer, S. et al. Diagnostic evaluation of river discharge into the Arctic Ocean and its impact on oceanic volume transports. Hydrology and Earth System Sciences 26, 279–304 (2022).

71. Huang, Y., Tagliabue, A. & Cassar, N. Data-Driven Modeling of Dissolved Iron in the Global Ocean. Frontiers in Marine Science 9, (2022).

72. Picheral, M. et al. The Underwater Vision Profiler 5: An advanced instrument for high spatial resolution studies of particle size spectra and zooplankton. Limnology and Oceanography: Methods 8, 462–473 (2010).

73. Picheral, M. et al. The Underwater Vision Profiler 6: an imaging sensor of particle size spectra and plankton, for autonomous and cabled platforms. Limnology and Oceanography: Methods 20, 115–129 (2022).

74. Gorsky, G. et al. Digital zooplankton image analysis using the ZooScan integrated system. Journal of Plankton Research 32, 285–303 (2010).

75. Dugenne, M., et al. First release of the Pelagic Size Structure database: Global datasets of marine size spectra obtained from plankton imaging devices. Earth System Science Data Discussions 1–41 (2023) doi:10.5194/essd-2023-479.

76. Kiko, R. et al. A global marine particle size distribution dataset obtained with the Underwater Vision Profiler 5. Earth System Science Data 14, 4315–4337 (2022).

77. Neeley, A., et al. Standards and practices for reporting plankton and other particle observations from images. (2021).

78. Drago, L., et al. Global Distribution of Zooplankton Biomass Estimated by In Situ Imaging and Machine Learning. Frontiers in Marine Science 9, (2022).

79. Boyer, T. P. et al. World Ocean Atlas 2018. NOAA National Centers for Envrionmental Information (2018).

80. Tagliabue, A. et al. How well do global ocean biogeochemistry models simulate dissolved iron distributions? Global Biogeochemical Cycles 30, 149–174 (2016).

81. Gledhill, M. & Buck, K. The Organic Complexation of Iron in the Marine Environment: A Review. Frontiers in Microbiology 3, (2012).

82. Aumont, O., Ethé, C., Tagliabue, A., Bopp, L. & Gehlen, M. PISCES-v2: an ocean biogeochemical model for carbon and ecosystem studies. Geoscientific Model Development 8, 2465–2513 (2015).

83. Kostadinov, T. S. et al. Ocean color algorithm for the retrieval of the particle size distribution and carbon-based phytoplankton size classes using a two-component coated-sphere backscattering model. Ocean Science 19, 703–727 (2023).

84. Chelton, D. B., Schlax, M. G. & Samelson, R. M. Global observations of nonlinear mesoscale eddies. Progress in Oceanography 91, 167–216 (2011).

85. Elith, J., Leathwick, J. R. & Hastie, T. A working guide to boosted regression trees. Journal of Animal Ecology 77, 802–813 (2008).

86. Steele, R. J. & Raftery, A. E. Performance of Bayesian Model Selection Criteria for Gaussian Mixture Models | University of Washington Department of Statistics. Frontiers of Statistical Decision Making and Bayesian Analysis 113–130 (2010).

